# Alt-RPL36 downregulates the PI3K-AKT-mTOR signaling pathway by interacting with TMEM24

**DOI:** 10.1101/2020.03.04.977314

**Authors:** Xiongwen Cao, Alexandra Khitun, Yang Luo, Zhenkun Na, Thitima Phoodokmai, Khomkrit Sappakhaw, Elizabeth Olatunji, Chayasith Uttamapinant, Sarah A. Slavoff

## Abstract

Thousands of previously unannotated small and alternative open reading frames (alt-ORFs) have recently been revealed in the human genome, and hundreds are now known to be required for cell proliferation. Many alt-ORFs are co-encoded with proteins of known function in multicistronic human genes, but the functions of only a handful are currently known in molecular detail. Using a proteomic strategy for discovery of unannotated short open reading frames in human cells, we report the detection of alt-RPL36, a 148-amino acid protein co-encoded with and overlapping human RPL36 (ribosomal protein L36). Alt-RPL36 partially localizes to the endoplasmic reticulum, where it interacts with TMEM24, which transports the phosphatidylinositol 4,5-bisphosphate [PI(4,5)P_2_] precursor phosphatidylinositol from the endoplasmic reticulum to the plasma membrane. Knock-out of alt-RPL36 in HEK 293T cells increases PI(4,5)P_2_ levels in the plasma membrane, upregulates the PI3K-AKT-mTOR signaling pathway, and increases cell size. Four serine residues of alt-RPL36 are phosphorylated, and mutation of these four serines to alanine abolishes interaction with TMEM24 and, consequently, abolishes alt-RPL36 effects on PI3K signaling and cell size. These results implicate alt-RPL36 as a novel regulator of PI(4,5)P_2_ synthesis upstream of the PI3K-AKT-mTOR signaling pathway. More broadly, these results show that the alt-RPL36 transcript can express two sequence-independent polypeptides from overlapping ORFs that regulate the same process – protein synthesis – via different molecular mechanisms (PI3K signaling and ribosome composition), expanding our knowledge of the mechanisms by which multicistronic human genes function.

## Introduction

Recent advances in genomic and proteomic technologies have revealed that mammalian genomes harbor thousands of previously unannotated small and alternative open reading frames (smORFs, <100 amino acids, and alt-ORFs, >100 amino acids, respectively)^1,2^. These genes previously escaped annotation not just because of their short length, but because, as a class, they exhibit low homology to proteins of known function, and are enriched for initiation at near-cognate non-AUG start codons (approximately 50%)^3,4^. Interestingly, while many smORFs and alt-ORFs lie in regions of RNA previously annotated as non-coding, up to 30% of the alt-ORFs identified by LC-MS/MS overlap an annotated protein-coding sequence in a different reading frame^2^. A rapidly increasing number of smORFs and alt-ORFs have been shown to play important roles in mammalian biology^5,6^. For example, NBDY regulates the mRNA decapping complex in human cells^7^, PIGBOS regulates the endoplasmic reticulum (ER) stress response^8^, and the MIEF1 microprotein regulates mitochondrial protein translation^9^. In mouse, AW112010 is a required component of the innate immune response^10^, and myoregulin interacts with SERCA calcium channels in muscle^11^. Recently, a genome-scale CRISPR screen revealed that over 500 smORFs regulate growth of human induced pluripotent stem cells^12^. These findings demonstrate that assignment of functions to small proteins represents a major opportunity to gain new insights into biology.

Currently, there is proteomic evidence for hundreds of uncharacterized human smORFs and alt-ORFs that are co-encoded with annotated proteins and initiate at non-AUG start codons. These findings are, in many cases, supported by ribosome profiling, conservation analyses, and *in silico* prediction of functional domains and secondary structure^13–15^. However, a majority of multicistronic and/or non-AUG-initiated alt-ORFs remain biochemically uncharacterized. Notable exceptions include the MIEF1 microprotein, which is encoded in a 5’UTR and regulates translation by mitoribosomes^9^, and Aw112010, which initiates at a near-cognate start codon and is required for mucosal immunity^10^.

Multiple human ribosomal proteins are encoded in complex genes that co-express two independent functional proteins. For example, ribosomal proteins L40 and S27A are synthesized as preproteins fused to ubiquitin, and S30 as a fusion to ubiquitin-like protein^16^. A prior proteogenomics study revealed translation of a sequence-independent alt-ORF co-encoded with human 60S ribosomal protein L36 (RPL36)^17^, which has nonetheless remained unannotated, likely due to a lack of information about its start codon and function. In this study, we provide molecular and proteomic evidence for its existence and define the complete coding sequence of this novel protein, which we term alt-RPL36. The human RPL36 gene is therefore dual coding via initiation of the alt-ORF at a near cognate start codon that overlaps the RPL36 nucleotide sequence in an alternative reading frame.

We further demonstrate that alt-RPL36 interacts with and regulates TMEM24 (transmembrane protein 24, alternatively C2CD2L, C2 Domain-Containing Protein 2-Like). TMEM24 is anchored to the ER membrane and mediates ER-plasma membrane (PM) contacts, transporting the PI(4,5)P_2_ precursor phosphatidylinositol from the ER to the PM^18^. This activity is required to replenish PI(4,5)P_2_ after its phosphorylation by phosphoinositide 3-kinase (PI3K) enzymes to generate phosphatidylinositol-3,4,5-trisphosphate (PI(3,4,5)P_3_) and downstream signaling lipids^19^, which activate the AKT-mTOR pathway to control cell growth and protein synthesis^20,21^. The localization of TMEM24 to ER-PM contacts is regulated by dynamic phosphorylation of the C-terminal region of TMEM24. Ca^2+^-stimulated phosphorylation causes TMEM24 dissociation from the PM, and dephosphorylation allows TMEM24 to re-associate with the PM^18^. TMEM24 is highly expressed in the brain and pancreatic islets^22^, and loss of TMEM24 in insulin-secreting cells leads to a defect in insulin release^18,23^.

In this study, we provide evidence for expression of alt-RPL36 from an upstream non-AUG start codon in human *RPL36* transcript variant 2, and show that it partially localizes to the ER in human cells using genomic knock-in tagging and unnatural amino acid mutagenesis labeling strategies. We identify and map four phosphorylation sites that are present at high stoichiometry in alt-RPL36, and show that phosphorylation is required for interaction with TMEM24. Finally, we engineered specific alt-RPL36 knockout and rescue cell lines to demonstrate that loss of alt-RPL36 promotes increased plasma membrane PI(4,5)P_2_ levels, as well as increased activation of the PI3K-AKT-mTOR pathway and increased cell size. We also demonstrate that the regulation of PI3K-AKT-mTOR pathway requires TMEM24. These results implicate alt-RPL36 as a novel upstream regulator of phospholipid transport and PI3K-AKT-mTOR signaling and demonstrate that overlapping alt-ORFs in the *RPL36* gene play related, but mechanistically distinct, biological roles in *trans*.

## Results

### A GUG-initiated alternative protein is translated from *RPL36* transcript variant 2

Using a previously reported proteogenomic strategy for unannotated small protein discovery^2^, we identified two tryptic peptides that mapped uniquely to an alternative reading frame of human *RPL36* transcript variant 2 in HEK 293T cells, which we name alt-RPL36 (Fig. 1a, b, and Supplementary Table 1). These tryptic fragments were also previously identified in the supplementary information of a proteogenomic study of A431 cells but were not characterized^17^. Alt-RPL36 tryptic peptides are also detected in HT1080 and MOLT4 cells (Supplementary Fig. 1a, b, and Supplementary Table 1), consistent with expression in several human cell lines from different tissues of origin. Compared to transcript variant 1 (NCBI RefSeq NM_033643), *RPL36* variant 2 (NCBI RefSeq NM_015414 and Fig. 1a) contains a longer 5’UTR, which we hypothesized to contain a start codon initiating alt-RPL36 translation. Interestingly, the first stop codon in frame with the observed tryptic peptides is downstream of the *RPL36* stop codon, meaning that alt-RPL36 is longer than RPL36 and completely encompasses its coding sequence. However, since alt-RPL36 is translated in the −1 reading frame relative to RPL36, the amino acid sequences of these two proteins are completely different (Fig. 1a).

**Fig. 1.**
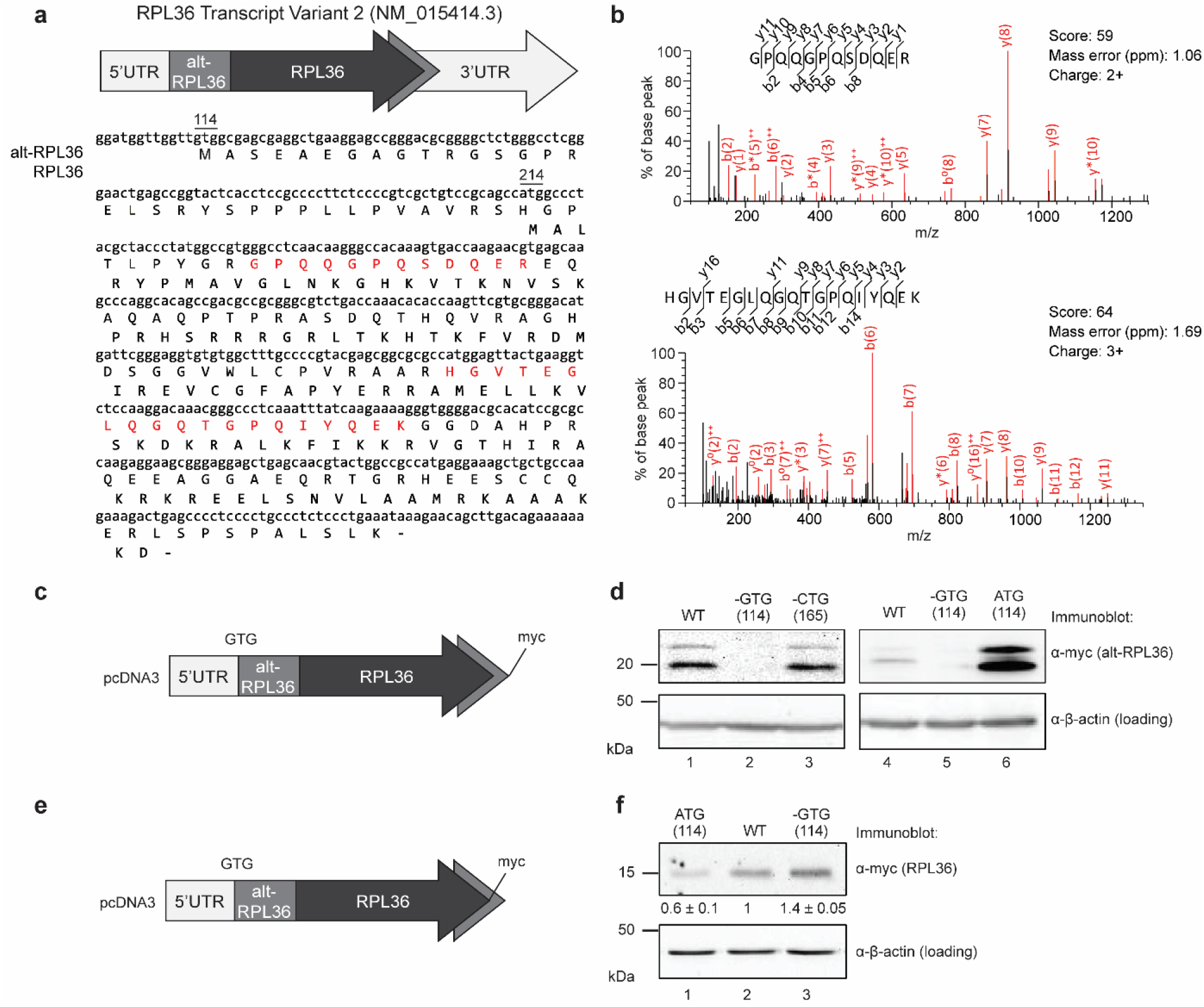
A GUG-initiated alternative protein is translated from *RPL36* transcript variant 2. **a** Top, a schematic representation of human RPL36 transcript variant 2 (tv2); light gray arrow, 5’ and 3’ untranslated regions (UTR); mid gray, alternative open reading frame (alt-ORF) encoding alt-RPL36; dark gray, annotated *RPL36* coding sequence. Bottom, the cDNA sequence of human *RPL36* transcript variant 2 is shown with the protein sequences of alt-RPL36 and RPL36 (bold) indicated below. The GTG start codon of alt-RPL36 and ATG start codon of RPL36 are numbered above the cDNA sequence. Highlighted in red are two tryptic peptides of alt-RPL36 detected by MS/MS. **b** MS/MS spectra of two alt-RPL36 tryptic peptides detected via peptidomics in HEK 293T cells. **c** and **d** Expression of a construct containing the full 5’UTR and alt-RPL36 coding sequence derived from *RPL36* tv 2, with a myc tag appended to the C-terminus of alt-RPL36 (**c**), in HEK 293T cells, was followed by lysis and Western blotting with the antibodies indicated to the right (**d**). **e** and **f** Expression of a construct containing the full 5’UTR and alt-RPL36 coding sequence derived from *RPL36* tv 2, with a myc tag appended to the C-terminus of RPL36 (**e**), in HEK 293T cells, was followed by lysis and Western blotting with the antibodies indicated to the right (**f**). Quantitative analysis of the Western blot signal of RPL36-myc are indicated at the bottom. Data represent mean values ± standard error of the mean (s.e.m.) of three biological replicates. All Western blots are representative of results obtained in three biological replicates.

To confirm expression and identify the start codon of alt-RPL36, the cDNA sequence comprising the 5’UTR of *RPL36* transcript variant 2 through the stop codon of the putative alt-ORF was cloned into a mammalian expression vector with a myc tag appended to the 3’ end of alt-RPL36. This construct produces two anti-myc immunoreactive bands (~22 kDa and ~20 kDa apparent molecular weight, due to phosphorylation, *vide infra*) from the alt-RPL36 reading frame when transiently transfected into HEK 293T cells (Fig. 1d, lanes 1 and 4). Because there is no upstream AUG start codon in frame with the observed alt-RPL36 tryptic peptides, we hypothesized that alt-RPL36 initiates at a near-cognate start codon, which can initiate protein translation with methionine at a fractional efficiency relative to AUG^24,25^. We searched the upstream, in-frame DNA sequence of alt-RPL36, and found two possible near-cognate start codons in a strong Kozak sequence context: G_114_TG and C_165_TG (numbered relative to the first nucleotide of the cDNA). Deletion of G_114_TG, but not C_165_TG, abolished the expression of alt-RPL36 (Fig. 1d, lanes 2, 3 and 5), and mutation of G_114_TG to A_114_TG increased the expression of alt-RPL36, consistent with increased efficiency of initiation at AUG codons^26^ (Fig. 1d, lane 6), indicating that G_114_TG is the start of the alt-RPL36 coding sequence.

Epitope-tagging of the RPL36 coding sequence revealed that the annotated ribosomal protein is also translated from *RPL36* transcript variant 2, and that translation of alt-RPL36 has a small (~40%) inhibitory effect on RPL36 synthesis from transcript variant 2. This is consistent with co-regulated expression of both ORFs from this transcript, as previously reported for uORFs initiating at non-AUG start codons^26^ (Fig. 1e, f). Taken together, these results indicate that human *RPL36* transcript variant 2 generates both RPL36 and an alternative protein, alt-RPL36, that initiates from G_114_TG, in overlapping reading frames.

To determine whether alt-RPL36 is conserved among species, RPL36 mRNAs from different species were obtained from NCBI nucleotide database, then translated in the +1, +2 and +3 frames using the ExPaSy translate tool. Cognate or near-cognate start codons within Kozak consensus motifs in frame with sequences homologous to human alt-RPL36 were identified in the 5’UTR of each transcript in order to predict the full-length sequence of hypothetical alt-RPL36 homologs. ClustalW alignment of hypothetical alt-RPL36 homologs from cat, cattle, and monkey revealed significant sequence similarity (Supplementary Fig. 2c), though protein-level evidence for these hypothetical homologs does not currently exist.

### Endogenously expressed alt-RPL36 partially localizes to the endoplasmic reticulum

Next, we wanted to determine whether alt-RPL36 is endogenously expressed from the *RPL36* genomic locus. To this end, we generated a knock-in (KI) HEK 293T cell line with a 3xGFP11-FLAG tag appended to the 3’ end of alt-RPL36. FLAG-IP followed by Western blotting revealed that the KI cells produced two specific anti-FLAG immunoreactive bands with the expected molecular weights, one major lower-mobility band and one faint higher mobility band, which were consistent with the over-expression results and which were absent in control HEK 293T cells, indicating the endogenous expression of alt-RPL36 (Fig. 2a).

**Fig. 2.**
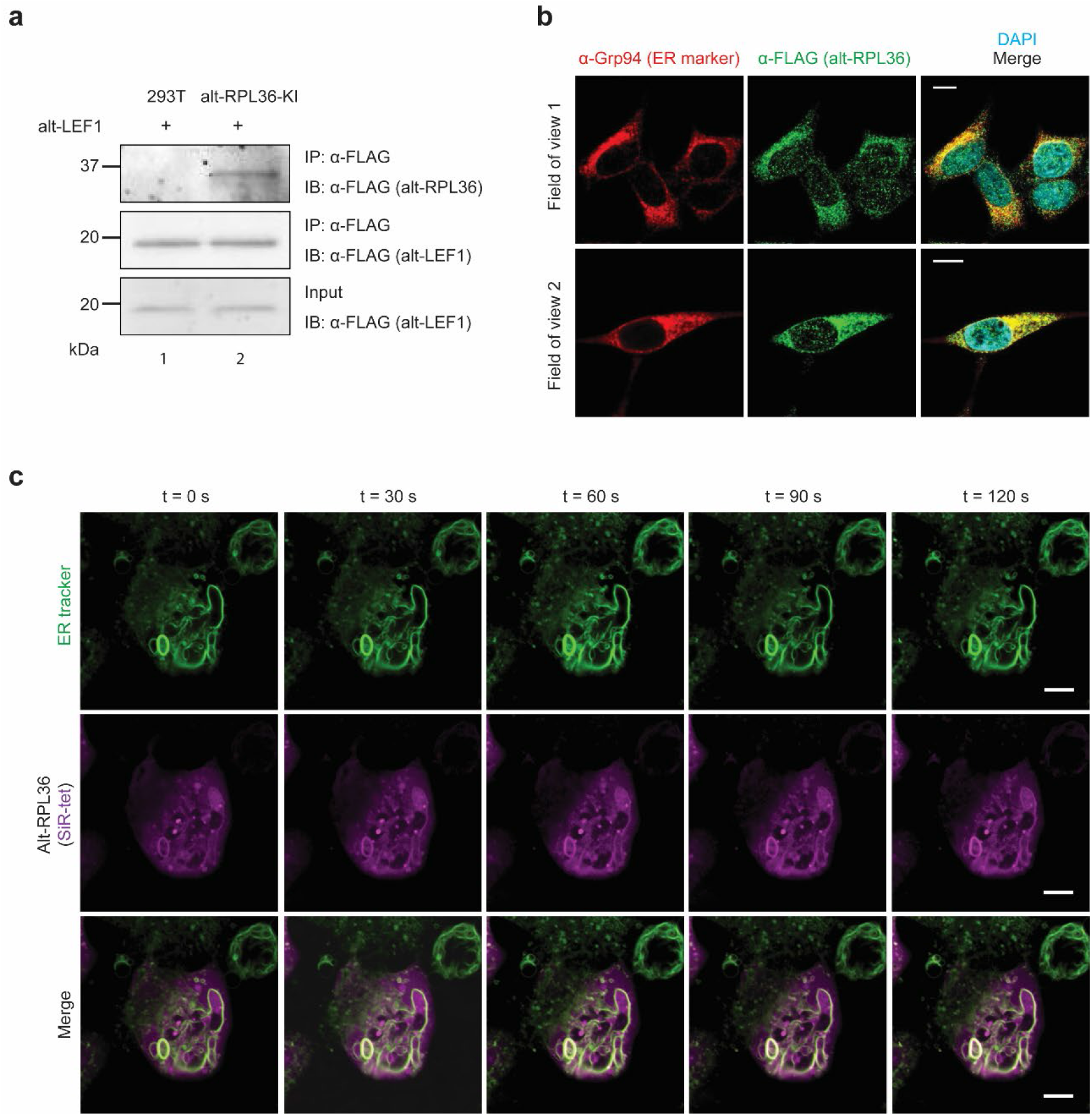
Alt-RPL36 is endogenously expressed and partially localizes to the endoplasmic reticulum. **a** Control HEK 293T cells (lane 1) or alt-RPL36-GFP11-FLAG knock-in (KI) HEK 293T cells (lane 2) were transfected with alt-LEF1-FLAG (which is a newly identified ~20 kDa alt-ORF^1^), serving as a FLAG-IP control, and IPs were performed with anti-FLAG antibody followed by IB with anti-FLAG. Cell lysates (1%) before IP (input) were used as the loading controls. **b** Confocal imaging of alt-RPL36-GFP11-FLAG KI cells after immunostaining with anti-Grp94 (red), antiFLAG (green) and DAPI (cyan). Scale bar 10 μm. **c** Live-cell imaging of alt-RPL36 via genetic code expansion. HEK 293T cells were transfected with plasmids expressing *M. mazei* pyrrolysyl-tRNA synthetase bearing Y306A/Y384F mutations (PylRS-AF), amber-suppressing Pyl tRNA (PylT_CUA_), and alt-RPL36^L18TAG^. Cells were incubated with 60 μM bicyclononyne-lysine (BCNK) for 45 hours, then labeled with tetrazine-silicon rhodamine (SiR-tet, magenta) and a live-cell ER tracker DPX dye (green) for 30 min before live-cell imaging. Time-lapse imaging over 2 min was shown. Scale bar 5 μm.

To further confirm the endogenous expression and investigate the subcellular localization of alt-RPL36, we performed immunostaining with the KI cells. As shown in Figure 2b, endogenously expressed alt-RPL36-3xGFP11-FLAG partially co-localizes with Grp94, which is an ER marker, indicating that a subset of alt-RPL36 localizes to ER.

To further confirm the subcellular localization of alt-RPL36, we visualized dynamics of alt-RPL36 in living cells via genetic code expansion-mediated labeling. Here a single amino acid residue of alt-RPL36 was replaced with a bicyclononyne-lysine (BCNK) unnatural amino acid via an engineered amber-suppressor pyrrolysyl tRNA (Pyl tRNA_CUA_/Pyl tRNA synthetase (PylRS) pair^27^. Alt-RPL36 bearing BCNK is subsequently derivatized with a membrane-permeable tetrazine-silicon rhodamine (SiR) conjugate via inverse-electron-demand Diels-Alder reaction for visualization in live cells^27^. We identified one amber variant—alt-RPL36^L18TAG^—which showed BCNK-dependent expression to produce full-length alt-RPL36 (Supplementary Fig. 3), and could be labeled in live HEK 293T cells with tetrazine-SiR. SiR-labeled alt-RPL36^L18BCNK^ was primarily cytosolic—which could be an artefact of overexpression under genetic code expansion—but a subpopulation of the protein showed clear overlap with ER tubules (Fig. 2c), suggesting ER-localization of alt-RPL36. Taken together, these results indicate that alt-RPL36 is endogenously expressed and partially localizes to the ER.

### Four serine residues of alt-RPL36 are phosphorylated

To test whether alt-RPL36 exhibits two bands by Western blot analysis (Fig. 1d, lane 1) due to protein phosphorylation^28^, alt-RPL36 was immunopurified from HEK 293T cells stably expressing alt-RPL36-FLAG-HA and treated with a nonspecific phosphatase. Phosphatase treatment eliminated the upper band, suggesting that it represents a phosphorylated form of alt-RPL36 (Fig. 3a). To identify the phosphorylated residues, we performed LC-MS/MS with immunopurified, digested alt-RPL36, and identified four candidate phosphoserine residues (S19, S22, S140 and S142) (Fig. 3b, Supplementary Table 2).

**Fig. 3.**
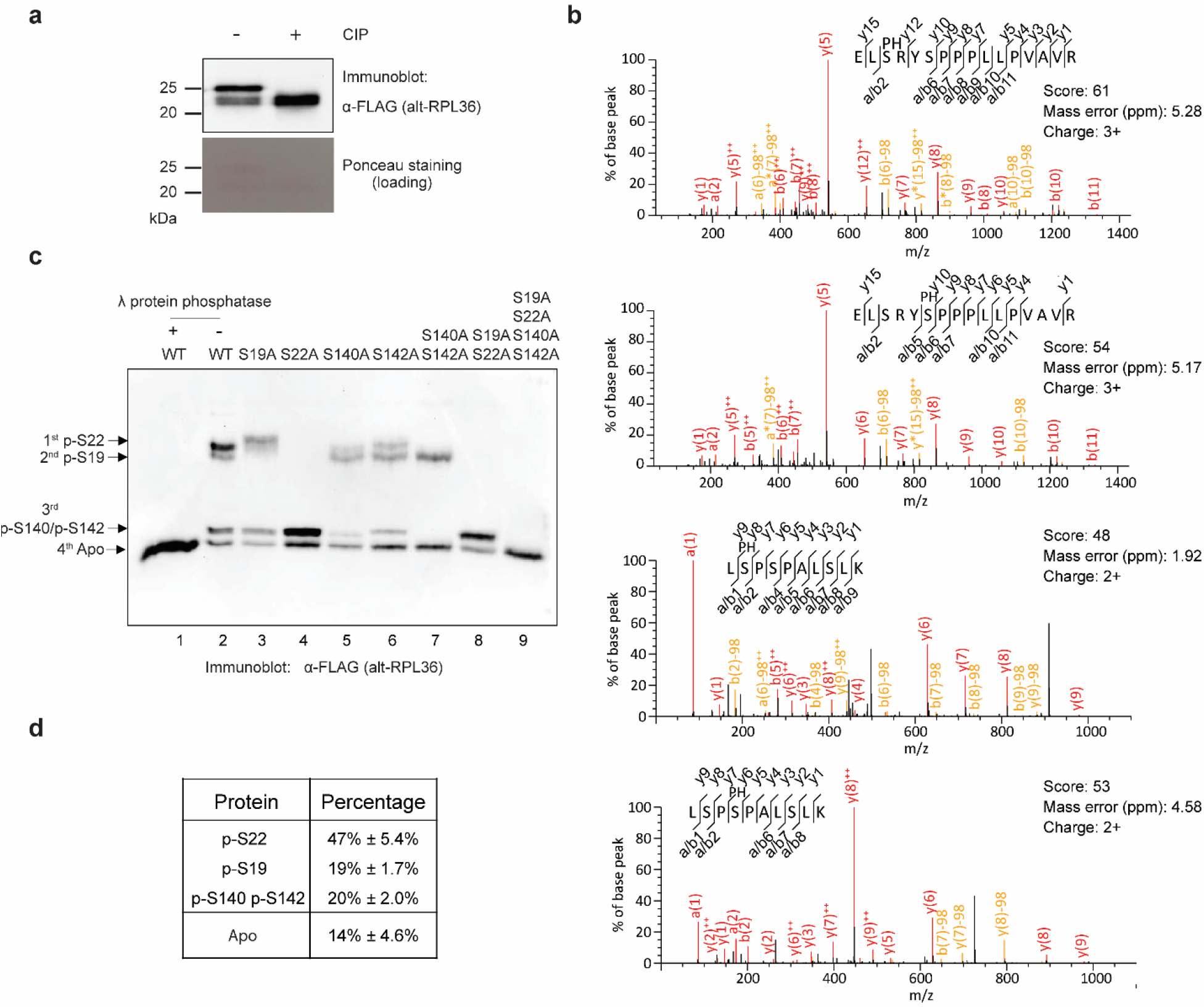
Four serine residues of alt-RPL36 are phosphorylated. **a** Western blot analysis of immunoprecipitated alt-RPL36-FLAG protein treated with calf intestinal alkaline phosphatase (CIP) and untreated control run on an SDS-PAGE gel. Ponceau staining served as a loading control. Data are representative of three biological replicates. **b** MS/MS spectra of alt-RPL36 phosphopeptides. S(PH) denotes phosphoserine. See Supplementary Table 2 for complete phosphoproteomics data. **c** To confirm phosphoserine assignments from MS/MS, wild-type alt-RPL36-FLAG or alanine point mutants were immunopurified from HEK 293T cells, resolved with Phos-tag SDS-PAGE, and detected with anti-FLAG antibody via Western blot. A sample of wild-type alt-RPL36-FLAG was treated with λ protein phosphatase before Phos-tag SDS-PAGE (lane 1). **d** Quantitative analysis of the Western blot signal of wild-type alt-RPL36-FLAG. Data represent mean values ± standard error of the mean (s.e.m.) of three biological replicates.

Because no phosphoantibodies specific to alt-RPL36 yet exist, we employed Phos-tag SDS-PAGE and Western blotting, combined with mutational analysis, to confirm the phosphorylation sites identified by LC-MS/MS. As shown in Figure 3c (lane 2), alt-RPL36 exhibited four bands corresponding to constitutively phosphorylated variants. Only the bottom band remained after nonspecific phosphatase treatment, indicating that it represents unphosphorylated alt-RPL36 (Fig. 3c, lane 1). Mutation of S19 to alanine abolished the second band, and mutation of S22 to alanine eliminated both the first and the second bands (Fig. 3c, lanes 3 and 4), suggesting that the second band represents pS19, the first band is pS22, and pS22 is required for phosphorylation of S19. Single mutation of S140 or S142 to alanine attenuated the signal of the third band (Fig. 3c, lanes 5 and 6), and double mutation of S140 and S142 to alanine abolished the third band entirely (Fig. 3c, lane 7), suggesting the third band represents both S140 and S142-phosphorylated alt-RPL36. The quadruple mutant S19A S20A S140A S142A exhibited a single band comigrating with phosphatase-treated wild-type alt-RPL36, further confirming that these four serine residues are the phosphorylation sites. Quantitation of replicate Western blots revealed that approximately 86% of alt-RPL36 is constitutively phosphorylated (Fig. 3d). High-occupancy phosphorylation of four specific sites in alt-RPL36 is consistent with a functional role for the protein.

### Phosphorylated alt-RPL36 interacts with TMEM24 via the SMP and C2 domains

Because many small proteins characterized to date bind to and regulate other proteins^29^, we performed a two-step co-immunoprecipitation (co-IP) of dually FLAG- and HA-tagged alt-RPL36 from HEK 293T cells and identified proteins specifically enriched over untransfected controls via quantitative proteomics. We then excluded common contaminants and proteins nonspecifically enriched by other previously reported, intrinsically disordered microproteins^7,30–32^. This analysis revealed that phospholipid transfer protein TMEM24/C2CD2L specifically co-immunopurifies with alt-RPL36 (Fig. 4a, Supplementary Table 3). To confirm the LC-MS/MS results, we performed reciprocal co-IP and Western blotting and observed enrichment of overexpressed alt-RPL36-myc by TMEM24-FLAG over controls (Fig. 4b, c).

**Fig. 4.**
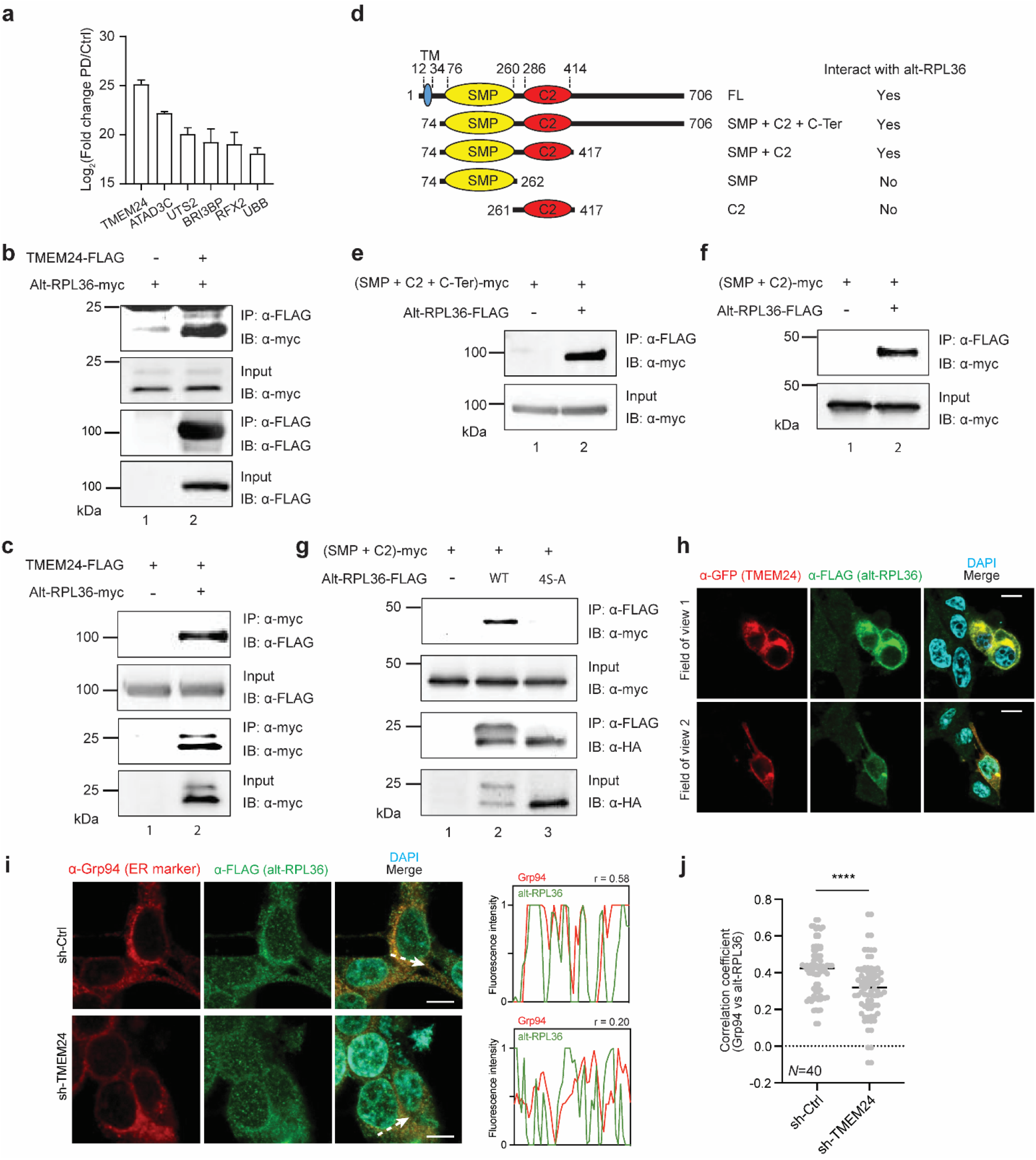
Phosphorylated alt-RPL36 interacts with TMEM24 via the SMP and C2 domains. **a** Quantitative proteomics (*N* = 2) of alt-RPL36-Flag-HA pulldown (PD) from HEK 293T lysates identified putative alt-RPL36 interaction partners enriched over untransfected HEK 293T cells (Ctrl). For complete quantitative proteomics results, see Supplementary Table 3. **b** and **c** HEK 293T cells were transfected with alt-RPL36-myc only (**b**, lane 1), TMEM24-FLAG only (**c**, lane 1) or both plasmids (**b** and **c**, lane 2), and immunoprecipitation (IP) was performed with anti-Flag (**b**) or anti-myc (**c**) antibody, followed by immunoblotting (IB) with antibodies indicated on the right. Cell lysates (1%) before IP (input) were used as the loading controls. **d** Schematic representation of the domain structures of the wild-type and truncation mutants of TMEM24, with amino acid residue numbers above. Alt-RPL36 interaction status of each construct is listed on the right. **e** and **f** Control HEK 293T cells (lane 1) or HEK 293T cells stably expressing alt-RPL36-FLAG-HA (lane 2) were transfected with TMEM24 truncation mutants, and IPs were performed with anti-FLAG antibody followed by IB with anti-myc. **g** Control HEK 293T cells (lane 1), HEK 293T cells stably expressing wild-type alt-RPL36-FLAG-HA (lane 2), and HEK 293T cells stably expressing non-phosphorylatable 4S-A mutant alt-RPL36-FLAG-HA (lane 3) were transfected with the TMEM24 SMP + C2 domain construct, followed by IP with anti-FLAG antibody and immunoblotting with antibodies indicated on the right. Cell lysates (1%) before IP (input) were used as the loading controls. All Western blots are representative of three biological replicates. **h** alt-RPL36-GFP11-FLAG KI cells were transfected with TMEM24-GFP, followed by immunostaining with anti-GFP (red), anti-FLAG (green) and DAPI (cyan). Scale bar 10 μm. **i** Confocal imaging of alt-RPL36-GFP11-FLAG KI cells stably expression control shRNA (sh-Ctrl, top panel) or TMEM24 shRNA (sh-TMEM24, bottom panel) after immunostaining with anti-Grp94 (red), anti-FLAG (green) and DAPI (cyan). Scale bar 10 μm. The arrow indicates the plane used for line profile generation. Right: line profiles of fluorescence intensities including Pearson correlation coefficients (r). **j** Pearson correlation coefficients of line profiles of Grp94 and alt-RPL36. *N*, number of line profiles. \*\*\*\**p*<0.0001.

TMEM24 consists of an N-terminal transmembrane domain that resides in the ER membrane, an SMP domain, a C2 domain, and an unstructured C-terminal region that interacts with the plasma membrane (Fig. 4d)^18^. To determine the domain(s) of TMEM24 with which alt-RPL36 interacts, we generated a series of truncation mutants of TMEM24 (Fig. 4d), transiently transfected these constructs into HEK 293T cells stably expressing alt-RPL36-FLAG-HA, and examined their ability to interact with alt-RPL36 by co-IP and Western blotting. Deletion of the N-terminal or the C-terminal regions of TMEM24 maintained the interaction (Fig. 4e, f), while further deletion of either the C2 domain or the SMP domain abolished the interaction (Supplementary Fig. 4a, b), indicating that both SMP and C2 domains are required for the interaction with alt-RPL36.

Since alt-RPL36 is constitutively phosphorylated, we asked whether these modifications regulate its interaction with TMEM24. As shown in Figure 4g, while wild-type, phosphorylated alt-RPL36 interacts with TMEM24, the quadruple alanine mutant, which abolishes all phosphorylation of alt-RPL36, eliminates the interaction. This may be due to reduction of the binding affinity to TMEM24 or change of the subcellular localization of alt-RPL36. These results indicate that the interaction between alt-RPL36 and the SMP and C2 domains of TMEM24 is specific and requires phosphorylation of alt-RPL36.

To determine whether endogenously expressed alt-RPL36 co-localizes with TMEM24 in cells, we transiently transfected TMEM24-GFP into alt-RPL36-FLAG KI cells, followed with immunostaining. As shown in Figure 4h, endogenous alt-RPL36 co-localizes with overexpressed TMEM24, consistent with an interaction between these proteins in cells. The plasma membrane localization of both TMEM24-GFP and alt-RPL36-FLAG may be due to increased ER-plasma membrane junction formation upon TMEM24 over-expression, as previously reported^22^. To determine whether TMEM24 is required for the ER localization of endogenous alt-RPL36, we knocked down TMEM24 with shRNA in the KI cells(Supplementary Fig. 5a, b), followed by immunostaining. As shown in Figures 4i and 4j, TMEM24 knockdown reduced the correlation coefficient between ER marker Grp94 and alt-RPL36, suggesting that interaction with TMEM24 is required for the ER localization of alt-RPL36.

We hypothesized that, since alt-RPL36 likely exists in complex with TMEM24 in cells, the same kinase may recognize and phosphorylate both proteins. The C-terminal region of TMEM24 has been reported to be phosphorylated by protein kinase C (PKC)^18^. Treatment of purified recombinant alt-RPL36 with PKC led to generation of lower-migration phosphorylated species in a Phos-tag Western blot (Supplementary Fig. 6a), indicating that PKC can phosphorylate alt-RPL36 *in vitro*. To determine whether PKC regulates the phosphorylation of alt-RPL36 in cells, we treated the HEK 293T cells stably expressing alt-RPL36-FLAG-HA with PKC inhibitor Bisindolylmaleimide II^33^, followed by FLAG-IP and Phos-tag Western blot. As shown in Supplementary Figure 6b and 6c, treatment with PKC inhibitor partially changes the phosphorylation pattern of alt-RPL36 in a dose-dependent manner, indicating PKC may, directly or indirectly, partially contribute to regulation of alt-RPL36 phosphorylation in cells.

It has been reported that phosphorylation of TMEM24 is regulated by the cytosolic calcium level^18^. We therefore asked whether the phosphorylation of alt-RPL36 is also dynamically regulated by changes in cytosolic calcium. To this end, we treated HEK 293T cells stably expressing alt-RPL36-FLAG-HA with thapsigargin, which elevates cytosolic calcium levels by blocking the ER calcium pump (SERCA)^34^, then measured phosphorylation of alt-RPL36 via Phos-tag Western blot. As shown in Supplementary Figures 6d and 6e, the phosphorylation pattern of alt-RPL36 changed within 15 min of thapsigargin treatment, indicating that phosphorylation of alt-RPL36 is dynamically regulated by cytosolic calcium.

### Phosphorylated alt-RPL36 regulates PI(4,5)P_2_ transport and the PI3K-AKT-mTOR signaling pathway

TMEM24 is an ER-anchored membrane protein that transports the PI(4,5)P_2_ precursor phosphatidylinositol to the plasma membrane (PM) at ER-PM contact sites via its SMP domain^18^. Phosphatidyinositol is the precursor to PI(4,5)P_2_, which is converted by PI3K to PI(3,4,5)P_3_ to regulate downstream AKT and mTOR signaling pathways^19–21^. To determine whether alt-RPL36 regulates TMEM24-dependent phenotypes, we generated an alt-RPL36-specific knock-out (KO) HEK 293T cell line using CRISPR-Cas9. The CRISPR/Cas9 strategy targeted 210 nucleotides surrounding the alt-RPL36 start codon without perturbing the RPL36 coding sequence (Supplementary Fig. 7a). Expression of the transcript variant 2-specific exon containing the alt-RPL36 start codon was undetectable via mRNA-seq and RT-qPCR in KO cells, but detectable in wild-type HEK 293T cells (Supplementary Fig. 7a, b). In the KO cells, expression of RPL36 was unchanged at the protein level, as indicated by Western blotting, demonstrating that specific deletion of the alt-RPL36 start codon did not affect the total RPL36 protein level (Supplementary Fig. 7c). To determine that any observed phenotypic effects are specific to alt-RPL36 ablation in the KO, and not off-target effects, we also generated a “rescue” cell line in which the alt-RPL36 coding sequence was stably reintroduced on the KO background, as well as “4S-A rescue” cells, in which the non-phosphorylatable mutant of alt-RPL36 was reintroduced on the KO background.

We hypothesized that the effect of alt-RPL36 on TMEM24-dependent phosphatidylinositol transport could have outcomes at multiple levels: (1) PI(4,5)P_2_ production in the plasma membrane immediately downstream of phosphatidylinositol transport, (2) AKT-mTOR activation downstream of PI(4,5)P_2_ phosphorylation by PI3K, and (3) transcriptional outputs downstream of AKT-mTOR pathway activation. Finally, phenotypic consequences should be exerted on cell morphology and size, which are regulated by PI3K and mTOR^35,36^. First, we assayed PI(4,5)P_2_ levels at the plasma membrane in wild-type, alt-RPL36 knock-out, rescue and 4S-A rescue HEK 293T cell lines stably expressing PH-PLC-GFP, which is a PI(4,5)P_2_ sensor^37^. As shown in Figure 5a-c, knock-out of alt-RPL36 increased the PH-PLC-GFP signal intensity compared with wild-type HEK 293T. The increase can be rescued by reintroduction of the coding sequence of wild-type, but not phosphorylation-incompetent, alt-RPL36. These results suggest that phosphorylated alt-RPL36 inhibits TMEM24-dependent phosphatidylinositol transport, therefore inhibiting production of PI(4,5)P_2_.

**Fig. 5.**
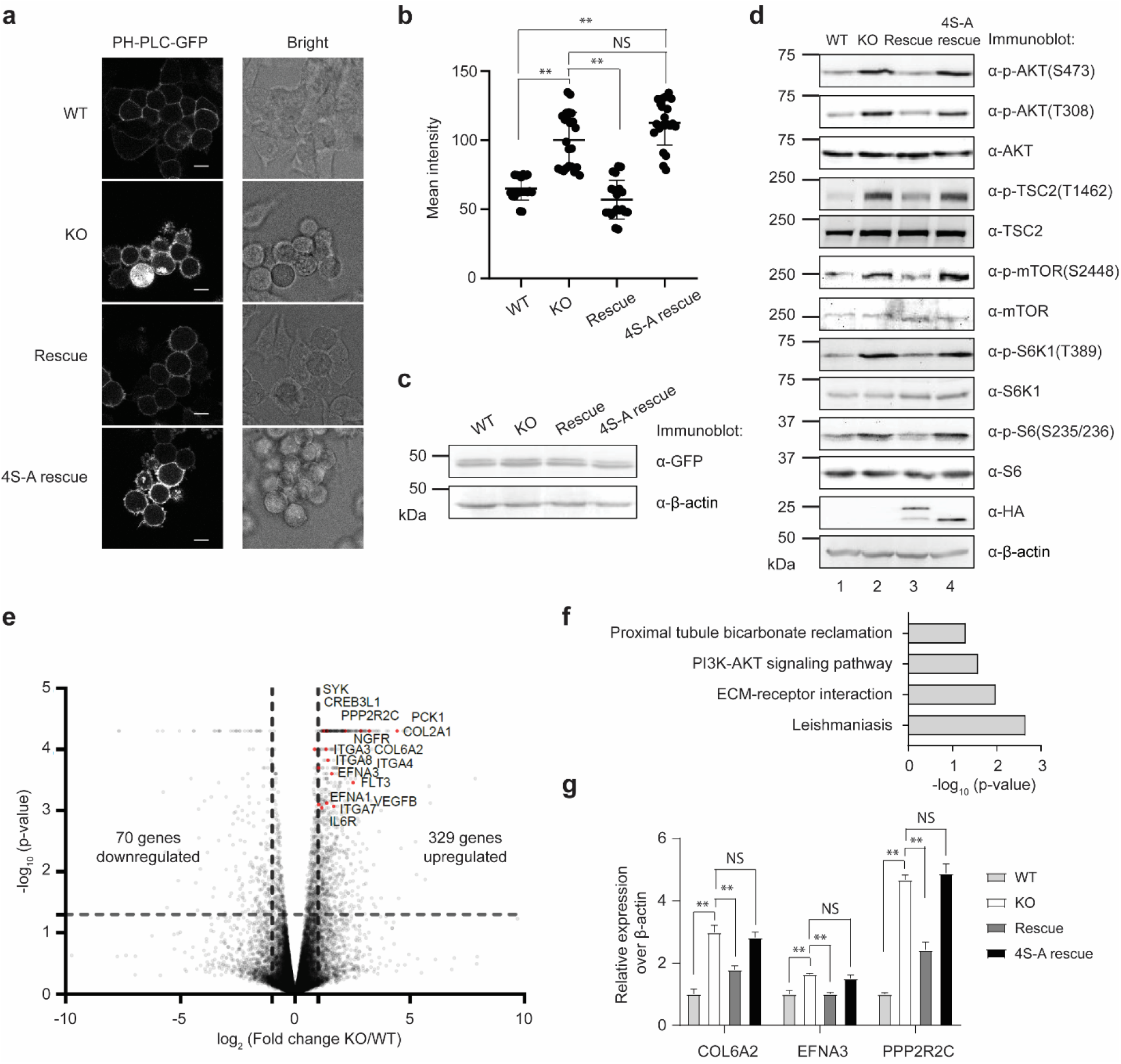
Phosphorylated alt-RPL36 regulates the PI3K-AKT-mTOR signaling pathway. **a** Confocal live-cell imaging of wild-type (WT), alt-RPL36 knock-out (KO), rescue with wild-type alt-RPL36 (Rescue) or rescue with nonphosphorylatable 4S-A mutant alt-RPL36 (4S-A rescue) HEK 293T cells stably expressing PH-PLC-GFP. Scale bar, 10 μm. **b** Quantitation of the PH-PLC-GFP signals in the four cell lines described above. At least 20 fields of view were analyzed, totaling > 400 cells for each measurement. Significance was evaluated with two-tailed *t*-test. ** *p* < 0.01. NS, not significant. **c** Western blot of the four cell lines with antibodies indicated on the right for comparison of PH-PLC-GFP expression. Data are representative of three biological replicates. **d** Western blot analysis of the wild-type (WT), alt-RPL36 knock-out (KO), rescue with wild-type alt-RPL36-FLAG-HA (Rescue) or rescue with 4S-A mutated alt-RPL36-FLAG-HA (4S-A rescue) HEK 293T cells with antibodies indicated on the right. Data are representative of three biological replicates. **e** Volcano plot of mRNA-seq from wild-type (WT) and alt-RPL36 knock-out (KO) HEK 293T cells. Upregulated transcriptional targets of the PI3K-AKT-mTOR signaling pathway identified via KEGG analysis are indicated in red and gene names are labeled. **f** KEGG analysis of genes upregulated upon alt-RPL36 knock-out; enriched biological pathways are plotted by significance (Fisher’s exact test). **g** Quantitative RT-PCR results of the mRNA transcripts of the wild-type (WT), alt-RPL36 knock-out (KO), rescue with wild-type alt-RPL36-FLAG-HA (Rescue) or rescue with 4S-A mutated alt-RPL36-FLAG-HA (4S-A rescue) HEK 293T cells. Data represent mean values ± standard error of the mean (s.e.m.) of four biological replicates. Significance was evaluated with two-tailed *t*-test. ** *p* < 0.01. NS, not significant.

To determine whether alt-RPL36 regulates the AKT-mTOR signaling pathway, we measured phosphorylation of these kinases and their substrates TSC2, S6K1 and S6, which correlates with pathway activation^20,21^. As demonstrated in Figure 5d, knock out of alt-RPL36 increased the phosphorylation of AKT, TSC2, mTOR, S6K1 and S6. Furthermore, this effect can be rescued by complementary expression of wild-type, but not phosphorylation-incompetent, alt-RPL36, consistent with the PI(4,5)P_2_ assay results. Interestingly, inhibition of mTOR by Torin1 reduced nascent alt-RPL36 translation, suggesting that mTOR regulates the translation of alt-RPL36 (Supplementary Fig. 8).

Third, we applied mRNA-seq to quantify the transcriptional state of wild-type vs. alt-RPL36 KO HEK 293T cells. We identified 329 genes significantly upregulated and 70 genes downregulated in the KO relative to wild-type (*p* < 0.05, *N* = 2, Fig. 5e). KEGG analysis of genes upregulated in the KO revealed that GO term “PI3K-AKT signaling pathway” is enriched (Fig. 5f). To validate the mRNA-seq data, we performed conventional RT-qPCR targeting three upregulated genes in the PI3K-AKT signaling pathway. As shown in Figure 5g, all three genes are upregulated upon alt-RPL36 knockout, compared with wild-type HEK 293T cells, consistent with the mRNA-seq results. The upregulation can be partially rescued by complementary expression of wild-type, but not phosphorylation-incompetent, alt-RPL36 (Fig. 5g). Taken together, these results suggest that phosphorylated alt-RPL36 downregulates the PI3K-AKT-mTOR signaling pathway.

Our results demonstrate that alt-RPL36 KO cells have an increased pool of PM PI(4,5)P_2_, which leads to a small basal increase in PI3K pathway signaling. This further suggests that cells should exhibit increased activation of PI3K signaling downstream of receptor tyrosine kinase signaling, including EGFR, which is a physiological activator of the PI3K signaling pathway^20^. Consistent with this notion, alt-RPL36 KO HEK 293T cells are more sensitive to EGF stimulation, indicated by increased phosphorylation of AKT and its substrate, TSC2, after EGF treatment, compared with control cells (Supplementary Fig. 9).

Inhibition of mTOR has been reported to reduce cell size^35,36^. Because alt-RPL36 activates the mTOR pathway, we asked whether loss of alt-RPL36 could increase cell size. To this end, we measured the cellular volume of four cell lines (WT, KO, Rescue and 4S-A rescue). As shown in Figures 6a and 6b, knockout of alt-RPL36 indeed increased cell size, indicated by a larger cell volume, compared with wild-type HEK 293T cells. The increased cell size can be rescued by wild-type, but not phosphorylation-incompetent, alt-RPL36. These results indicate that alt-RPL36 downregulates cell size, consistent with inhibition of TMEM24 and the PI3K-AKT-mTOR pathway.

**Fig. 6.**
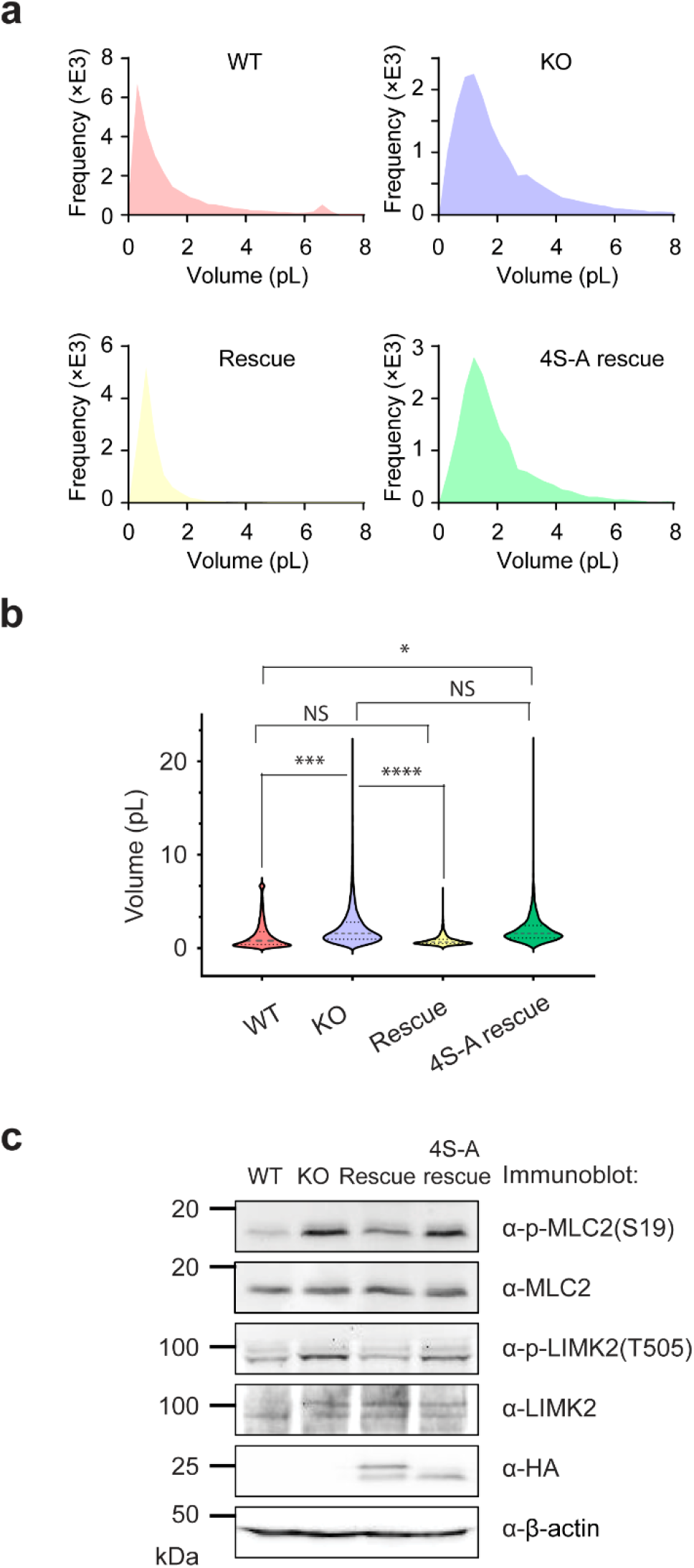
Alt-RPL36 regulates cell size and morphology. **a** Cell volume histogram for wild-type (WT), alt-RPL36 knock-out (KO), rescue with wild-type alt-RPL36 (Rescue) or rescue with nonphosphorylatable 4S-A mutant alt-RPL36 (4S-A rescue) HEK 293T cells. **b** Violin plot of the cell volume of the four cell lines described above. Significance was evaluated with non-parametric Mann-Whitney test. * *p* < 0.05, *** *p* < 0.001, **** *p* < 0.0001. NS, not significant. **c** Western blot of the four cell lines indicated at the top with the antibodies indicated to the right. Data are representative of three biological replicates.

We noticed that the KO and 4S-A rescue HEK 293T cells showed a round morphology (Fig. 5a, rows 2 and 4), though this effect is only partially rescued by reintroduction of alt-RPL36 (Fig. 5a, row 3). We hypothesized that this could be caused by PI3K-dependent activation of the ROCK1 signaling pathway^38–40^, which induces cytoskeletal reorganization to promote cell motility. To confirm this hypothesis, we measured the phosphorylation levels of two ROCK1 substrates, MLC2 and LIMK2. As shown in Figure 6c, the phosphorylation of MLC2 and LIMK2 are upregulated upon alt-RPL36 knockout, and the upregulation can be partially rescued by wild-type, but not phosphorylation-incompetent, alt-RPL36. The observed rounding of the rescue cells may be due to incomplete rescue by alt-RPL36 reintroduction. Taken together, these results indicate that alt-RPL36 regulates cell morphology via the ROCK1 pathway.

### Regulation of PI3K-AKT-mTOR pathway by alt-RPL36 requires its interaction with TMEM24

To determine whether regulation of PI3K-AKT-mTOR pathway by alt-RPL36 requires its interaction with TMEM24, we knocked down TMEM24 with shRNA on the background of alt-RPL36 knockout (Fig. 7c, d), then measured PM PI(4,5)P_2_ levels and PI3K-AKT-mTOR pathway phosphorylation. As shown in Figures 7a and 7b, knock out of alt-RPL36 increased PI(4,5)P_2_ levels compared to wild-type HEK 293T when transfected with a control shRNA (sh-Ctrl), as expected. Knockdown of TMEM24 on the alt-RPL36 KO background partially rescued the increase of PI(4,5)P_2_ compared to transfection of alt-RPL36 KO cells with sh-Ctrl. These results indicate that alt-RPL36-dependent regulation of PI(4,5)P_2_ levels requires TMEM24. Similarly, as shown in Figure 7d, while knock-out of alt-RPL36 increased phosphorylation levels of PI3K-AKT-mTOR pathway kinases and substrates compared to wild-type (compare lane 2 with lane 1), knockdown of TMEM24 in alt-RPL36 KO partially rescued the increases (compare lane 4 with lane 2), again indicating that alt-RPL36-dependent regulation of PI3K-AKT-mTOR pathway requires TMEM24.

**Fig. 7.**
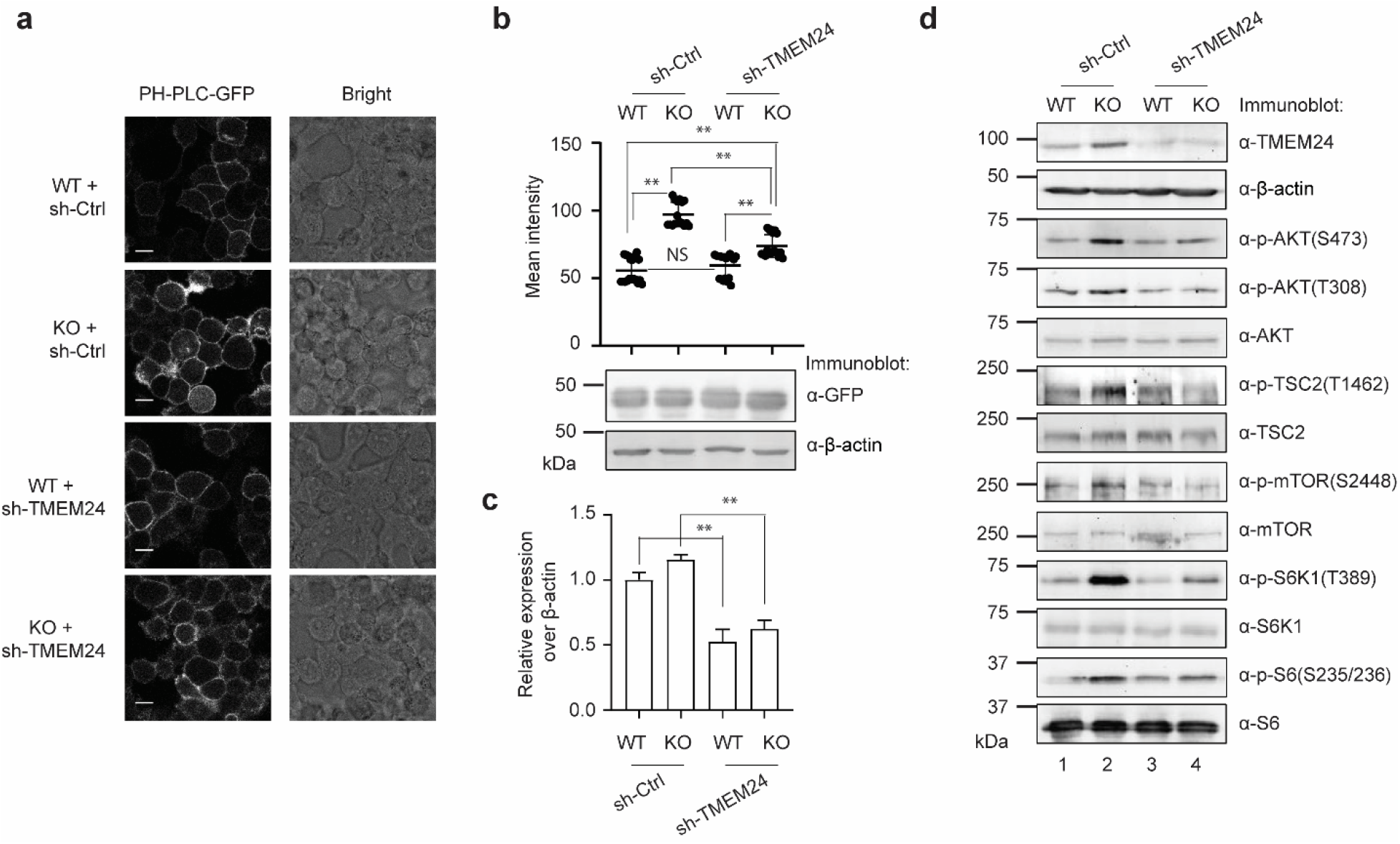
Regulation of PI3K-AKT-mTOR pathway by alt-RPL36 requires its interaction with TMEM24. **a** PH-PLC-GFP reporter was stably introduced into the following cell lines: HEK 293T cells stably expressing control shRNA (WT + sh-Ctrl), alt-RPL36 KO cells stably expressing control shRNA (KO + sh-Ctrl), HEK 293T stably expressing TMEM24 shRNA (WT + sh-TMEM24), alt-RPL36 KO stably expressing TMEM24 shRNA (KO + sh-TMEM24). Confocal imaging was performed with live cells. Scale bar, 10 μm. **b** Top: quantitation of the PH-PLC-GFP signals in the four cell lines described above. At least 15 fields of view were analyzed, totaling > 300 cells for each measurement. Significance was evaluated with two-tailed *t*-test. ** *p* < 0.01. NS, not significant. Bottom: Western blot of the four cell lines with antibodies indicated on the right for comparison of PH-PLC-GFP expression. Data are representative of three biological replicates. **c** Quantitative RT-PCR with primers specific to TMEM24 to assess efficiency of silencing by sh-TMEM24 (error bars, standard error of the mean (s.e.m.), *N* = 4, ** *p* < 0.01 (two-tailed *t*-test)). **d** Western blot analysis of the four cell lines described above with antibodies indicated on the right. Data are representative of three biological replicates.

To further confirm these results, we knocked out TMEM24 in alt-RPL36 KO cells by transfecting them with Cas9 lentivirus and a guide RNA targeting TMEM24. We then measured PM PI(4,5)P_2_ levels and the phosphorylation levels of the kinases and substrates of the PI3K-AKT-mTOR pathway, and observed similar results (Supplementary Fig. 10). Taken together, these results confirm that the regulation of PI3K-AKT-mTOR pathway by alt-RPL36 requires its interaction with TMEM24.

It is important to note that knock down of TMEM24 in wild-type HEK 293T cells did not significantly change PM PI(4,5)P_2_ levels (Fig. 7a and Supplementary Fig. 9a, compare row 3 with row 1) or activation of the PI3K-AKT-mTOR pathway (Fig. 7d and Supplementary Fig. 9d, compare lane 3 with lane 1). We hypothesize that this is because TMEM24 is nearly completely inhibited by alt-RPL36 under basal conditions in cultured HEK 293T cells.

### Cancer-related mutations affect alt-RPL36 protein properties and interaction

A number of cancer-associated point mutations in the human *RPL36* gene have been previously reported, and we queried whether any of these mutations could alter the sequence of alt-RPL36 (Supplementary Fig. 11a). Of 10 cancer-specific point mutations in the (alt-)RPL36 coding sequence (CDS) in the COSMIC database (cancer.sanger.ac.uk)^41^, 2 alter the amino acid sequence of RPL36 but are synonymous in the alt-RPL36 reading frame; 4 affect both the RPL36 and alt-RPL36 amino acid sequences; and, remarkably, 4 are synonymous with respect to RPL36 but create nonsynonymous mutations in the alt-RPL36 reading frame.

To determine whether any of these mutations affect the properties or function of alt-RPL36, we generated HEK 293T stable cell lines expressing seven cancer-associated alt-RPL36 mutants. We first examined their effect on protein expression. As shown in Supplementary Figure 11b, four point mutations (V93A, S50N, Q98H and P115A) exhibited reduced protein expression levels relative to the wild-type alt-RPL36 sequence. The mRNA profile of these mutants did not match the protein levels, suggesting that altered protein expression is likely post-transcriptionally regulated (Supplementary Fig. 11c). Second, we examined the phosphorylation state of these mutants via Phos-tag Western blotting. As shown in Supplementary Figure 11d, 5 of the 7 mutants (V93A, S50N, Q98H, Q104H and P115A) changed the phosphorylation pattern of alt-RPL36. The E118K mutation changed the mobility of all alt-RPL36 protein bands in the Phos-tag Western blot, likely by changing the overall charge state of the protein. Finally, we tested the ability of each alt-RPL36 mutant to interact with TMEM24. As shown in Supplementary Figure 11e, the S50N mutation essentially completely abolished the interaction, and other four mutations (R73C, Q98H, Q104H and P115A) reduced the interaction. Taken together, these results suggest that several cancer-associated alt-RPL36 variants exhibit altered expression and phosphorylation, which may affect alt-RPL36 function. In the case of the melanoma-associated S50N variant, the observed loss-of-function with respect to TMEM24 interaction would be expected to derepress TMEM24.

## Discussion

In this work, we have identified a previously unannotated protein, alt-RPL36, that is co-encoded with human ribosomal protein L36 in the −1 reading frame of *RPL36* transcript variant 2. Alt-RPL36 initiates at a non-AUG start codon upstream of the RPL36 initiation site and terminates downstream of the RPL36 stop codon, and both of these sequence-independent proteins are expressed from the same mRNA. Previous reports have demonstrated that upstream ORFs with non-AUG start codons tend to exhibit positively correlated expression regulation with downstream coding sequences^42^, while upstream AUG start codons strongly inhibit translation of downstream cistrons^43^. Therefore, because alt-RPL36 initiates at a non-AUG start site, it does not efficiently repress downstream translation of RPL36 from the downstream AUG start codon, allowing the two proteins to be translated from the same mRNA. Despite their co-expression, alt-RPL36 is likely present in cells at a significantly lower concentration than RPL36, though we have not yet quantified its cellular copy number. The transcript variant 2-specific exon is ~10-fold less abundant than an exon specific to transcript variant 1, which only encodes RPL36, according to quantitative RNA-seq (Supplementary Fig. 7a). Further taking into account ~10-fold less efficient translation initiation at GUG start codons relative to AUG^44^, there is likely to be at least 100-fold less alt-RPL36 in the cell than RPL36. Notwithstanding their differing cellular abundances, RPL36 – a ribosomal protein – and alt-RPL36 – a negative regulator of the AKT-mTOR pathway – regulate the same process (protein translation) via different mechanisms, and in different directions. While many human smORFs are now hypothesized to exist in bicistronic transcripts that function as operons^12,14^, few examples of co-encoded human proteins have been characterized in molecular detail. This includes bicistronically-encoded fatty acid synthesis and RNA processing proteins HsHTD2 and RPP14, and overlapping open reading frames encoding RNA-binding proteins and interactors ATXN1/Alt-ATXN1^45,46^. This work, combined with previous literature, expands the recent finding that dense co-encoding of functionally related proteins within the same transcript - and even within overlapping nucleotide sequences - can occur in the human genome^45–48^.

Because the alt-RPL36 and RPL36 coding sequences overlap, conservation of alt-RPL36 may be challenging to detect. Furthermore, it is not yet clear whether alternative splice variants of *RPL36* exist, or are annotated, in different species. Expression of putative homologs in other mammalian species must be experimentally tested. It is also possible that alt-RPL36 represents an example of a *de novo* evolved gene^49,50^. Functional *de novo* evolved genes have been previously detected in plants and animals, wherein they function in reproduction and development and are differentially expressed in response to stress^51,52^.

Previous proteomic studies have detected phosphorylated peptides mapping to alt-proteins in cells undergoing mitosis and stimulation by EGF^13,53^. This data strongly suggests that alt-ORFs may be functional and regulated by post-translational modifications. The discovery that alt-RPL36 is highly phosphorylated, and that its association with its target protein requires these phosphoserine residues, further supports the notion that additional sm/alt-ORF-encoded proteins could bear functional modifications. Similar to TMEM24^18,22^, phosphorylation of alt-RPL36 is also dynamically regulated by the cytosolic calcium level. While we have not yet explored the functional consequence of this regulation, considered together with the observation that alt-RPL36 phosphorylation is required for TMEM24 association, these results suggest that repression of TMEM24 by phosphorylated alt-RPL36 could potentially be reversible under some physiological conditions.

The discovery of alt-RPL36 provides new opportunities for investigating dynamic upstream regulation of the PI3K-AKT-mTOR pathway, an overall model of which is provided in Figure 8. Phosphorylated (P) alt-RPL36 interacts with and negatively regulates TMEM24-dependent phosphatidylinositol (PI) transport. In cells lacking alt-RPL36, or, hypothetically, cells in which alt-RPL36 phosphorylation is reduced, TMEM24 is released from inhibition by alt-RPL36, increasing PI transport to the plasma membrane and subsequent conversion to PI(4,5)P_2_ and PI(3,4,5)P_3_ – the latter via the activity of PI3K. PI(3,4,5)P_3_ activates AKT, leading to activation of downstream mTOR signaling and upregulates protein translation – likely including translation of alt-RPL36. The resulting increase in alt-RPL36 then inhibits TMEM24, restoring homeostasis.

**Fig. 8.**
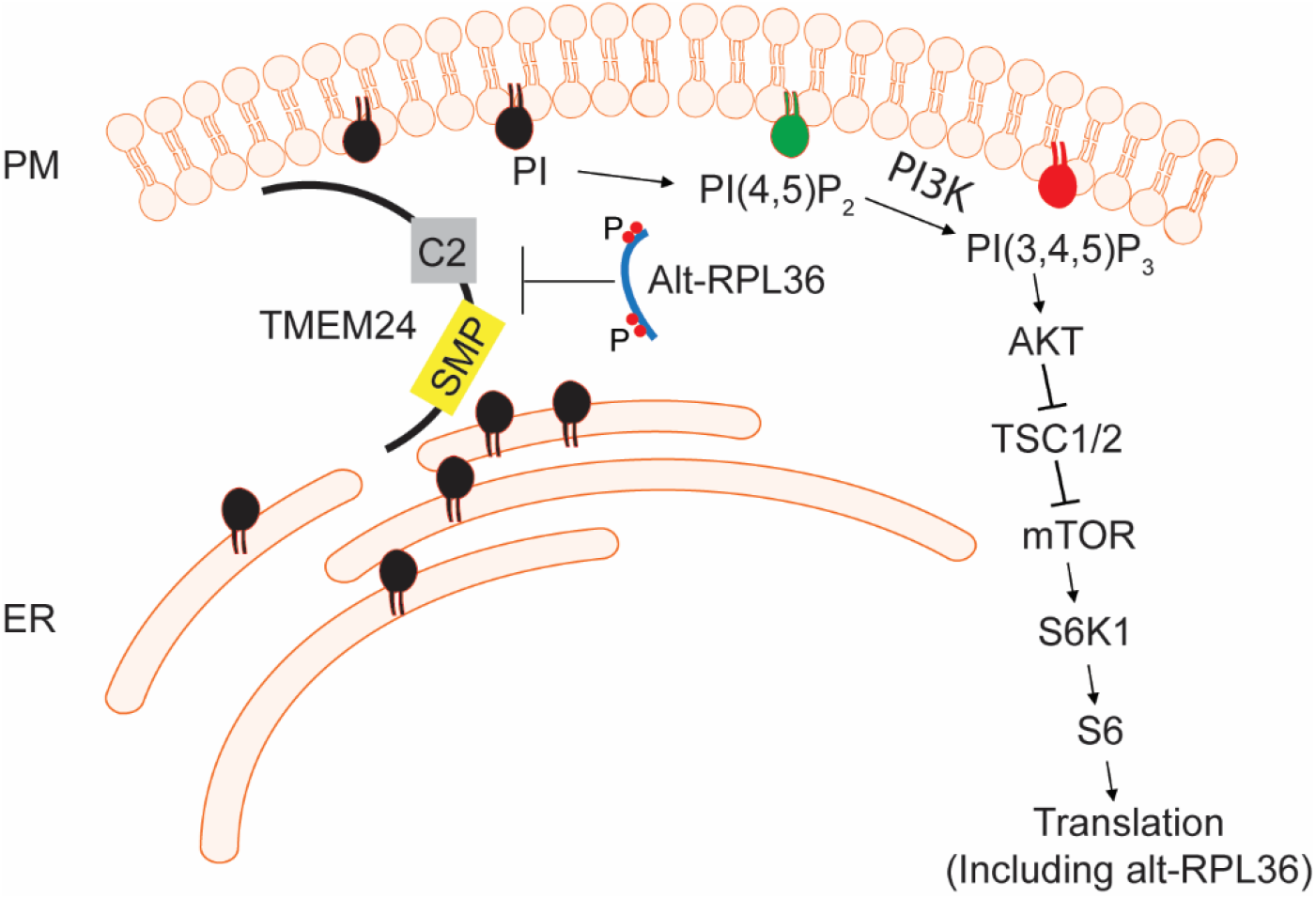
Model of alt-RPL36 regulatory pathway. Phosphorylated (P) alt-RPL36 interacts with TMEM24 via the SMP and C2 domains to negatively regulate TMEM24-dependent phosphatidylinositol 4,5-bisphosphate [PI(4,5)P_2_] precursor phosphatidylinositol (PI) transport. In cells lacking alt-RPL36, TMEM24 is released from the inhibition by alt-RPL36, and more PI is transported to the plasma membrane (PM) from the endoplasmic reticulum (ER), where it is synthesized. At the PM, PI is converted to PI(4,5)P_2_ then phosphorylated by PI3K to produce PI(3,4,5)P_3_. PI(3,4,5)P_3_ then activates AKT, which phosphorylates and inhibits TSC1/2, then activates downstream mTOR signaling. mTOR phosphorylates S6K1, and activated S6K1 phosphorylates S6, finally upregulates protein translation, including alt-RPL36. Upregulated alt-RPL36 then inhibits TMEM24, restoring homeostasis.

The subcellular localization of alt-RPL36 may be relevant to its function, but several questions remain to be answered. TMEM24 has been reported to localize to ER-PM contact sites, and to relocalize throughout the ER when its C-terminal domain is phosphorylated. The observation that endogenously expressed alt-RPL36 localizes to the pan-ER, but is enriched at PM contact sites when over-expressing TMEM24, is consistent with TMEM24 binding. However, further work is needed to determine whether, like TMEM24, alt-RPL36 relocalizes under physiological conditions, and whether its subcellular localization is regulated by its phosphorylation state under physiological conditions.

Finally, our functional data demonstrate that the cellular phenotypes associated with alt-RPL36 ablation require TMEM24, conclusively establishing TMEM24 as the direct functional target of alt-RPL36. These data furthermore suggest that alt-RPL36 inhibits TMEM24. However, the functional assays employed in this study are indirect readouts of TMEM24 activity. Therefore, the mechanism by which phosphorylated alt-RPL36 inhibits TMEM24 remains to be established. It is possible that the alt-RPL36 interaction with the SMP and C2 domains of TMEM24 affects its lipid transport activity, either by competitive binding or conformational change. It is alternatively possible that phosphorylated alt-RPL36 contributes to electrostatic inhibition of the TMEM24-PM interaction, thereby interfering with phospholipid transport between these membranes by altering TMEM24 localization. A similar electrostatic inhibitory effect leading to PM dissociation has been shown to occur upon phosphorylation of the TMEM24 C-terminus^18,22^. Simpler mechanisms, such as alteration of the stability or folding of TMEM24 by alt-RPL36, are also possible. Cell-based, biochemical and structural studies will be required to differentiate these possible mechanisms of action.

## Online methods

### Data analysis

Statistics, two-tailed *t*-test was performed using Excel or Prism, and equal variance between samples being compared was established using an *F*-test.

### Antibodies and reagents

Primary antibodies for Western blotting include the following anti-FLAG (Sigma, F3165); anti-myc (Rockland, 600-401-381, Cell Signaling, 2276 (mouse), or Cell Signaling, 2278 (rabbit)); anti-HA (Invitrogen, 71-5500); anti-His (Thermo Fisher, MA1-21315); anti-V5 (Cell Signaling, 13202); anti-β-actin (Invitrogen, BA3R); anti-p-mTOR(S2448) (Cell Signaling Technology, 2971); anti-mTOR (Cell Signaling Technology, 2972); anti-p-AKT(S473) (Cell Signaling Technology, 9271); anti-p-AKT(T308) (Cell Signaling Technology, 4056); anti-AKT (Cell Signaling Technology, 9272); anti-p-TSC2(T1462) (Cell Signaling Technology, 3617); anti-TSC2 (Cell Signaling Technology, 4308); anti-p-S6K1(T389) (Cell Signaling Technology, 9234); anti-S6K1 (Cell Signaling Technology, 9202); anti-p-S6(S235/236) (Cell Signaling Technology, 2211); anti-S6 (Cell Signaling Technology, 2217); anti-p-MLC2(S19) (Cell Signaling Technology, 3671); anti-MLC2 (Cell Signaling Technology, 3672); anti-p-LIMK2(T505) (Cell Signaling Technology, 3841); anti-LIMK2 (Cell Signaling Technology, 3845); anti-p-EGFR(Y1068) (Abcam, ab40815); anti-EGFR (Abcam, ab52894); anti-RPL36 (Bethyl Laboratories, A305065A-M); anti-TMEM24 (a gift from Pietro De Camilli, Yale). Immunoprecipitation was performed with the following antibody beads: anti-FLAG M2 affinity gel (Sigma, A2220); anti-myc tag agarose beads (Sigma, A7470); anti-HA tag magnetic beads (Thermo Fisher, 88836). mTOR inhibitor Torin1 (Tocris, 4247), PKC inhibitor Bisindolylmaleimide II (Cayman, 11020), Tharpsigargin (Sigma, T9033), L-Azidohomoalanine (Click Chemistry tools, 1066-1000), Biotin-alkyne (Click Chemistry tools, 1266-5). Human EGF protein (Abcam, ab9697).

### Cloning and genetic constructs

A construct comprising the full 5’UTR of human *RPL36* transcript variant 2 through the stop codon of alt-RPL36 was synthesized by Genscript with a myc epitope tag appended to the 3’ end of the alt-RPL36 coding sequence, then subcloned into pcDNA3. For generation of HEK 293T cells stably expressing alt-RPL36, a dual FLAG and HA tag were appended to the 3’ end of alt-RPL36 by PCR, and the GTG start codon was mutated to ATG. The dually tagged coding sequence was amplified by PCR, then cloned into pLJM1. The serine point mutations with a dual FLAG and HA tag were generated by ligating the PCR products into AgeI and EcoRI cloning sites in the pLJM1 vector. The full-length TMEM24 clone with a C-terminal FLAG epitope tag in pcDNA3 was purchased from Genscript. The truncations of TMEM24 were generated by PCR with a C-terminal myc epitope tag, then subcloned into pcDNA3. The TMEM24-GFP plasmid is a gift from Dr. Pietro De Camilli at Yale. The cDNA clone expressing PH-PLC-GFP was purchased from Addgene (a gift from Tobias Meyer, Stanford), then cloned into pLJM1 by PCR. pPB-MmPylRS-AF-4xPylT_CUA_, which expresses *M. mazei* pyrrolysyl tRNA synthetase^Y306A/Y384F^ (PylRS-AF) and 4 copies of pyl tRNA_CUA_, was a gift from Jason Chin (MRC Laboratory of Molecular Biology, UK). pPB-alt-RPL36-4xPylT_CUA_ was cloned via subcloning alt-RPL36 gene to replace MmPylRS-AF via restriction cloning. Site-directed mutagenesis to introduce L18TAG, A30TAG, and V31TAG mutations to alt-RPL36 were performed on pPB-alt-RPL36-4xPylT with standard protocols. Constructs expressing subcellular APEX2--pcDNA5-V5-APEX2-NES (cytosolic targeting), pcDNA5-V5-APEX2-NLS (nuclear targeting), pcDNA5-V5-APEX2-Sec61β (targeting to cytosolic face of ER membrane), and pcDNA3-Sec61β-V5-APEX2 (ER lumen targeting) were gifts from Hyun-Woo Rhee (Seoul National University, South Korea). pcDNA5-alt-RPL36-APEX2 and pcDNA3-APEX2-alt-RPL36 were cloned via subcloning alt-RPL36 to replace Sec61β portion of pcDNA5-V5-APEX2-Sec61β and pcDNA3-Sec61β-V5-APEX2 respectively, via restriction cloning.

### Cell culture and transfection

HEK 293T cells were purchased from ATCC and early-passage stocks were established in order to ensure cell line identity; cells were maintained up to only 10 passages. HEK 293T cells were cultured in DMEM (Corning, 10-013-CV) with 10% FBS (Sigma, F0392) and 1% penicillin-streptomycin (VWR, 97063-708) in a 5% CO_2_ atmosphere at 37°C. Plasmid transfection was performed with Lipofectamine 2000 or Lipofectamine 3000 and Opti-MEM (GIBCO, 31985-070) according to the manufacturer’s instructions, or polyethyleneimine (PEI, Polysciences, 23966-1) according to established protocol^54^.

### Lentivirus production and stable cell line generation

Lentivirus was produced as previously described^55^. Briefly, HEK 293T cells were co-transfected using Lipofectamine 2000 or polyethyleneimine with expression construct in pLJM1, along with pMD2.G and psPAX2, and growth media were replaced after 3-5 h (Lipofectamine 2000 transfection) or 7-8 h (polyethyleneimine transfection). 48 h post-transfection, media containing viruses was harvested, filtered through a 0.45-μm filter, and infection was performed by mixing with two volume fresh media containing suspended HEK 293T cells. 24 h post-infection, the growth media was replaced. 48 h post-infection, stably expressing cells were selected with 6 μg/mL puromycin for 2 days. Early stocks of stable cell lines were established after selection. Stable cell lines were released from puromycin for 2 days prior to use in experiments.

### Alt-RPL36 labeling and imaging

HEK 293 cells were seeded onto 6-well plates for Western blot analysis or on glass coverslips placed in 24-well plates for imaging studies. The coverslips were incubated for 1 hour with 0.1 mg/mL poly-D-lysine. Upon reaching ~ 80% confluency, the media was exchanged for Opti-MEM containing transfection mixture of DNA (plasmid ratio of pPB-alt-RPL36-4xPylT_CUA_: pPB-mMPylRS-AF-4xPylT_CUA_ = 9: 1), Lipofectamine 3000 and its accessory reagent (Thermo Fisher). After incubation for 6 hours at 37 °C, the media was exchanged for DMEM supplemented with 10% FBS, 1% Pen-strep and 60 uM BCNK (Sichem). Subsequently, cells were grown in presence of BCNK for 45 hours. To analyze BCNK-bearing alt-RPL36 expression, cells were lysed with RIPA buffer, and lysates resolved on polyacrylamide gels. Proteins were transferred onto nitrocellulose membranes, and Western blotting with anti-myc antibody was performed using standard protocols and imaged with ImageQuant LAS500 (GE Healthcare).

To fluorescently label alt-RPL36 via inverse-electron-demand Diels-Alder cycloaddition, cells were grown in BCNK-free media for another 3-6 hours to remove excess BCNK. Thereafter, 800 nM tetrazine-SiR (Spirochrome) was incubated with the cells in regular growth media at 37° C for 30 minutes. To stain the ER, 0.5 μM of ER tracker Blue-White DPX dye (Thermo Fisher) was added simultaneously with tetrazine-SiR. Cells were washed extensively with PBS prior to livecell imaging on an Olympus Fluoview FV3000 confocal microscope with 63× oil immersion objective under atmosphere-controlled stage at 37°C and appropriate laser and filter settings, and analyzed with cellSens Software.

To determine whether alt-RPL36 is co-localized with ER marker Grp94, line profile analysis was performed. Line profiles were generated with FIJI (ImageJ). For each cell, two straight lines were drawn to cross the ER in different directions, indicated by the arrows shown in the figures. Fluorescence signals along the straight line of ER and examined proteins were calculated with the plot profile tool in FIJI. The Pearson’s correlation coefficient r values of two fluorescence signals were calculated with Excel. Perfect co-localization is indicated by r = 1, perfect exclusion is indicated by r = −1, and 0 means random distribution.

### PH-PLC-GFP imaging

HEK 293T cells stably expressing PH-PLC-GFP were grown to 80% confluency on coverslips in 12-well plates. Coverslips were inverted and imaged in pre-warmed DMEM with 10% FBS, 1% penicillin-streptomycin in MatTek imaging dishes. Confocal imaging was performed on a Leica SP8 LS confocal microscope with 63× oil immersion objective under atmosphere-controlled stage at 37°C. Quantification was performed with Image J using standard parameters.

### Co-immunoprecipitation

HEK 293T cells were grown to 50% confluency in 15 cm dishes, then transiently transfected with the indicated plasmids. 48 h after transfection, cells were harvested and suspended in 1 mL lysis buffer (Tris-buffered saline (TBS) with 1% Triton X-100 and Roche Complete protease inhibitor cocktail tablets). Then cells were sonicated (50% intensity, 5 s pulse with 25 s rest, 5×, MICROSON XL 2000) on ice followed by centrifugation at 15,000 r.p.m., 4°C, 10 min. 1% of lysate samples were saved for analysis of loading. A 25 μL aliquot of anti-FLAG agarose beads (clone M2, Sigma) was washed with 1 mL lysis buffer, collected by centrifugation for 2 min at 1,000 g, then suspended in the cell lysate supernatant. Beads suspensions were rotated at 4°C for 1 h, then washed 3 times with wash buffer (Tris-buffered saline (TBS) with 1% Triton X-100 and 350 mM NaCl), then 1 time with lysis buffer. Proteins were eluted by adding 40 μL SDS-PAGE loading buffer and boiling.

### Immunoprecipitation and proteomics

Control HEK 293T cells or HEK 293T cells stably expressing alt-RPL36-FLAG-HA were grown to 80-90% confluency in 15 cm dishes. Cells were harvested and immunoprecipitation were performed as described above. After the final wash, elution was in 40 μL of 3× FLAG peptide (Sigma), at a final concentration of 100 μg/mL in lysis buffer at 4°C for 1 h. Beads were removed by centrifugation and the entire supernatant were collected. A 50 μL aliquot of anti-HA magnetic beads (Cat.88836, Thermo Fisher) was washed with 1 mL lysis buffer, then suspended in the elution. Beads suspensions were rotated at 4°C for 4 h, then washed 3 times with lysis buffer. Elution was in 40 μL of HA peptide (Covance), at a final concentration of 400 μg/mL in lysis buffer at 4°C for 2 h. The eluted proteins were subjected to MS-MS analysis.

### Proteomics and database searches

Protein-containing gel slices were digested with trypsin at 37°C for 14-16 h. The resulting peptide mixtures were extracted from the gel, dried, followed with ethyl acetate extraction to remove residual detergent, then re-suspended in 15 μl of 3:8 70% formic acid:0.1% TFA. A 5 μL aliquot of each sample was injected onto a pre-packed column attached to a nanoAcquity UPLC (Waters) in-line with an LTQ Orbitrap Velos (Thermo Scientific) and a 90-min gradient was used to further separate the peptide mixtures as follows (solvent A: 0.1% formic acid; solvent B: acetonitrile with 0.1% formic acid): Single pump trapping was turned on for 6 min at a flow rate of 2.5 μL/min at 98% A. Isocratic flow was maintained at 0.3 μL/min at 2% B for 10 min, followed by linear gradients from 2% B to 10% B over 2 min, 10% B to 25% B over 58 min, 25% B to 40% B over 10 min, 40% B to 95% B over 2 min. Isocratic flow at 95% B was maintained for 5 min, followed by a gradient from 95% B to 2% B over 10 min. The column flow rate was 0.3 μL/min. The full MS was collected over the mass range of 298-1,750 m/z with a resolution of 30,000. MS/MS data was collected using a top 10 high-collisional energy dissociation method in data-dependent mode with a normalized collision energy of 33.0 eV and a 2.0 m/z isolation window. The first mass was 100 m/z in fixed mode. MS/MS resolution was 7,500 and dynamic exclusion was 60 seconds.

For identification of alt- and microproteins, ProteoWizard MS Convert was used for peak picking and files were analyzed using Mascot. Oxidation of methionine and N-terminal acetylation were set as variable modifications, and a three-frame translation of mRNA-seq from HEK 293T cells was used as the database, as previously reported^1^. For co-IP proteomics searches and quantitative analysis, files were analyzed using MaxQuant, oxidation of methionine and N-terminal acetylation were set as variable modifications, and human UniProt plus alt-RPL36 was used as the database for searching. For phosphoproteomics searches, phosphorylation of Ser, Thr and Tyr, oxidation of methionine and N-terminal acetylation were set as variable modifications. ProteoWizard MS Convert was used for peak picking and files were analyzed using Mascot, and human UniProt plus alt-RPL36 was used as the database. For all analysis, a mass deviation of 20 p.p.m. was set for MS1 peaks, and 0.6 Da was set as maximum allowed MS/MS peaks with a maximum of two missed cleavages. Maximum false discovery rates (FDR) were set to 1% both on peptide and protein levels. Minimum required peptide length was five amino acids. Protein quantitation was accomplished via spectral counting, where the number of total peptides observed for each identified protein was taken as the total spectral counts and compared for the IP vs. negative control samples.

### Phos-tag SDS PAGE

Phos-tag SDS-PAGE was performed with 10% polyacrylamide gels containing 50 μM Phos-tag acrylamide and 100 μM MnCl2. After electrophoresis, Phos-tag acrylamide gels were washed with transfer buffer containing 10 mM EDTA for 10 min with gentle shaking and then with transfer buffer without EDTA for 10 min according to the manufacturer’s protocol. Proteins were transferred to nitrocellulse membranes followed by a standard Western blotting protocol^7^.

### Phosphatase treatment

Alt-RPL36-FLAG-HA expressed and immunopurified from HEK 293T cells was incubated with 0.4 U/ μL alkaline phosphatase, calf intestinal (CIP) (NEB, cat. M0290) at 37°C for 1 h, or 8 U/ μL lambda protein phosphatase (NEB, cat. P0753) at 30°C for 0.5 h according to the manufacturer’s instructions in a total reaction volume of 20 μL before Western blotting.

### Recombinant expression and purification of human alt-RPL36

His-MBP-tagged human alt-RPL36 in pET21a was transformed into an *E. coli*. BL21 (DE3) strain. Overnight cultures were diluted 1:500 in Luria-Bertani (LB) media supplemented with 100 μg/mL of ampicillin and grown at 37°C. Expression was induced by adding 0.5 mM of isopropyl β-d-1-thiogalactopyranoside (IPTG) when OD 600 reached 0.6, followed by shaking at 18°C overnight. After cell harvest and resuspension in lysis buffer (50 mM Tris-HCl pH 7.5, 300 mM NaCl, 2 mM imidazole) with 1 mM β-mercaptoethanol and Roche Complete protease inhibitor cocktail tablets, cells were lysed by sonication (30% intensity, 10 s pulse with 50 s rest on ice, 20×). The solution was clarified by centrifugation at 14,000 rpm for 20 min at 4°C and loaded onto a column containing 1 mL of Co^2+^-TALON resin (Takara, 635606) pre-equilibrated with lysis buffer. Following incubation for 1 h at 4°C, the resin was washed with 10 mL of lysis buffer and 30 mL wash buffer (50 mM Tris-HCl pH 7.5, 300 mM NaCl and 10 mM imidazole), and proteins were eluted with elution buffer (50 mM Tris-HCl pH 7.5, 300 mM NaCl and 250 mM imidazole). Eluted proteins mixed with 15% glycerol were snap-frozen and stored at −80°C.

### In vitro phosphorylation

In vitro alt-RPL36 phosphorylation by PKCα (Invitrogen, P2232) was performed by incubating various concentrations (0-0.3 μM) of PKCα with 2 μM purified His-MBP-alt-RPL36 in reaction buffer (20 mM HEPES-KOH pH 7.4, 16.7 mM CaCl2, 10 mM DTT, 100 mM MaCl2, 100 μM ATP and 0.6 mg/mL phosphatidylserine) at 30°C for 0.5 h in a total reaction volume of 10 μL.

### Generation of alt-RPL36 knock out and knock in cell lines

Alt-RPL36 KO and 3xGFP11-FLAG KI HEK 293T cells were generated using CRISPR-Cas9. Guide RNAs (gRNAs) were designed with the guide design tool from the Zhang lab (crispr.mit.edu) to target the RPL36 genomic region (gRNA1: 5’-CCGGGATATCTACTCGGCTC-3’; gRNA2: 5’-GAGTACCGGCTCAGTTCCCG-3’) for KO, and gRNA 5’-CCCGATAGTCGCCGTCTCGG-3’ for KI. Double-stranded DNA oligonucleotides corresponding to the gRNAs were inserted into pSpCas9(BB)-2A-GFP vector (Addgene, as a gift from F. Zhang, MIT, Cambridge, MA). For generation of KO cells, an equal mixture of the two gRNA plasmids were transfected into HEK 293T cells using Lipofectamine 2000 (Invitrogen) according to the manufacturer’s instructions, and GFP-positive cells were sorted with flow cytometry. Loss of alt-RPL36 expression was confirmed by genomic DNA PCR and sequencing. In the alt-RPL36 KO cell line used in this study, the two alleles were disrupted by a 210-nt homozygous deletion, deleting the alt-RPL36 start codon, but not the RPL36 start codon. And no large deletion or insertion were observed in the 100 kb upstream and 100 kb downstream of the edited sites, indicated by Xdrop long-read DNA sequencing (Supplementary Figure 12 and Table 4). Eleven predicted off-target sites were confirmed to be unedited by genomic PCR and sequencing (Supplementary Figure 13 and Table 5).

For generation of KI cells, a donor plasmid containing 300 bp homology left-arm and 300 bp homology right-arm sequence around the stop codon of alt-RPL36, which are separated with 3xGFP11-FLAG tag and BamHI / NotI restriction sites were synthesized from GenScript, a DNA sequence containing pGK promoter and hygromycin resistant gene were subcloned into the donor plasmid with BamH1 and NotI. An equal mixture of the gRNA and donor plasmids were transfected into HEK 293T cells using Lipofectamine 2000 (Invitrogen) according to the manufacturer’s instructions, and hygromycin selection were performed 2-day post transfection. Alt-RPL36-3xGFP11-FLAG KI cells were confirmed by genomic DNA PCR and sequencing. And no large deletion or insertion were observed in the 100 kb upstream and 100 kb downstream of the edited sites, indicated by Xdrop long-read DNA sequencing (Supplementary Figure 12 and Table 4).

### KO and KI cell line validation through Xdrop long-read DNA sequencing

High molecular weight (HMW) DNA was isolated from alt-RPL36 knock-out (KO) or knock-in (KI) cell lines using the Blood & Cell Culture DNA Mini Kit (Qiagen). Xdrop sequencing were performed by Samplix (Denmark). Briefly, HMW DNA were evaluated by the TapestationTM System (Agilent Technologies Inc.), using Genomic DNA ScreenTape according to the manufacturer’s instructions. The HMW DNA samples were further purified using HighPrep™ PCR Clean-up Bead System according to the manufacturer’s instructions (MAGBIO Genomics) with the following changes: Bead-to-sample ratios were 1:1 (v:v) and elution was performed by heating the sample in elution buffer for 3 minutes at 55 °C before separation on the magnet. The samples were eluted in 20 μl 10 mM Tris-HCl (pH 8). Purified DNA samples were quantified by Quantus (Promega Inc.) FluorometerTM, according to the manufacturer’s instructions. DNA was partitioned in droplets by Xdrop™ and subjected to PCR using the enrichment PCR assays. The primers were designed ~45 kb both up- and downstream of RPL36 on chromosome 19 with the aim of enriching 100 kb of DNA in either direction. Primers were localized to genes CATSPERD and SAFB at positions chr19:5,720,156-5,778,734 and chr19:5,623,035-5,668,478, respectively. The droplet productions were then sorted by fluorescence activated cell sorting (FACS). The isolated droplets were broken, and DNA was again partitioned in droplets by Xdrop™ and amplified by droplet Multiple Displacement Amplification (dMDA) reactions. After amplification DNA was isolated and quantified. Minion Oxford Nanopore Sequencing platform was used to generate long-read sequencing data from the dMDA samples as described by the manufacturer’s instructions (Premium whole genome amplification protocol (SQK-LSK109) with the Native Barcoding Expansion 1-12 (EXP-NBD104)). Generated raw data (FAST5) was subjected to basecalling using Guppy v.3.4.5 with high accuracy and quality filtering to generate FASTQ sequencing data. A minimum of 1 gigabyte of sequencing data was obtained from each sample. Raw reads were corrected using the necat.pl script from the package NECAT (v 0.0.1). Corrected reads were further split at chimeric positions using the SACRA pipeline (https://github.com/hattori-lab/SACRA) according to standard parameters. Finally non-chimeric reads were mapped to GRCh38 chromosome 19 with ngmlr 0.2.7 and variants were called using sniffles 1.0.12 using parameters specific to ONT sequencing data^56^.

### Alt-RPL36 conservation analysis

For alt-RPL36 conservation analysis, RPL36 mRNAs from different mammalian species were obtained from the NCBI nucleotide database, then translated in the +1, +2 and +3 frames using ExPaSy translate tool. Cognate or near-cognate start codons with in-frame Kozak consensus motifs were identified in the 5’UTR of each transcript, and were considered as the first amino acid (methionine) of alt-RPL36. The hypothetical proteins thus derived were aligned with Clustal Omega and visualized using JalView software using standard parameters.

### Cell size measurements

The Scepter handheld cell counter (Millipore) was used to perform cell diameter and volume measurements on cells grown to 80-90% confluency and resuspended in PBS to a concentration of 1-5 10^5^ cells/mL. Measurements were taken using a 60 μM sensor. Data were exported and visualized using Scepter Software Pro 2.1. Cell size distributions were compared for statistical differences using the non-parametric Mann-Whitney test.

### mRNA-seq and data analysis

Whole RNA was isolated from 2.5 x 10^6^ control or alt-RPL36 knockout (*N* = 2) HEK 293T cells using Qiagen RNeasy Mini Kit spin columns, then treated in solution with DNase I prior to Qiagen column clean-up according to the manufacturer’s protocol. Whole RNA was submitted to the Yale Center for Genomic Analysis for preparation according to the standard Illumina protocol for paired-end sequencing with enrichment of poly-A RNA. Samples were multiplexed and 75 bp fragments were sequenced on the Hiseq2500 sequencer. The reads were mapped to the human genome (hg19) using TopHat (v2.0.11). To identify the genes regulated by alt-RPL36, we counted the RNA reads in exons and calculated the reads per kilobase per million reads (RPKM) for each gene as a measure of expression using Cufflinks (v.2.2.1) with the Cuffdiff tool using default parameters. A log2 (fold change) cutoff of 0.2 with *p* value ≤ 0.05 was used to identify the genes with downregulated or upregulated expression in the alt-RPL36 knockout cells, compared with that in the control cells. Gene ontology analysis was performed using g:Profiler with standard parameters^57^. For qRT-PCR, total RNA was isolated using Trizol (Invitrogen) according to the manufacturer’s protocol. Reverse transcription was performed with iScript (Bio-Rad) and qPCR was performed with iTaq Universal SYBR Green Supermix (Bio-Rad), with quantitation by a relative Ct method. qPCR primer sequences are provided in Supplementary Table 6.

## Data availability

The mRNA-seq and Xdrop sequence data have been deposited in the NCBI Gene Expression Omnibus under accession GSE144979. Proteomics data were deposited under accession PXD018268 to the PRIDE repository.

## Author Contributions

X.C., A.K., Y. L. and E.O. designed and performed experiments and analyzed data. Z.N. designed and constructed CRISPR/Cas9 knockout cells. T.P. and K.S. performed imaging and APEX fingerprinting. C.U. designed experiments and analyzed data. S.A.S. conceived the project, designed experiments, and analyzed data. X.C., C.U., and S.A.S. wrote the manuscript, and all authors edited and approved the final version of the manuscript.

## Acknowledgment

We thank Benjamin Turk, Pietro de Camilli, and the members of the Slavoff lab for helpful discussions. No competing financial interests have been declared. This work was supported by an American Cancer Society Institutional Research Grant Individual Award for New Investigators (IRG-58-012-57), the Searle Scholars Program, an Odyssey Award from the Richard and Susan Smith Family Foundation, and Yale University West Campus start-up funds (to S.A.S.). X.C. was supported in part by a Rudolph J. Anderson postdoctoral fellowship from Yale University. A.K. was in part supported by an NIH Predoctoral Training Grant (5T32GM06754 3-12). T.P., K.S., and C.U. were supported by student and research assistant funds from VISTEC (to T.P. and K.S. respectively), a startup grant from VISTEC (to C.U.), Thailand Research Fund (MRG6280177, to C.U.), and Seed Awards in Science from Wellcome Trust (to C.U.).

