## Supplementary Information for "Alt-RPL36 downregulates the PI3K-AKT-mTOR signaling pathway by interacting with TMEM24"

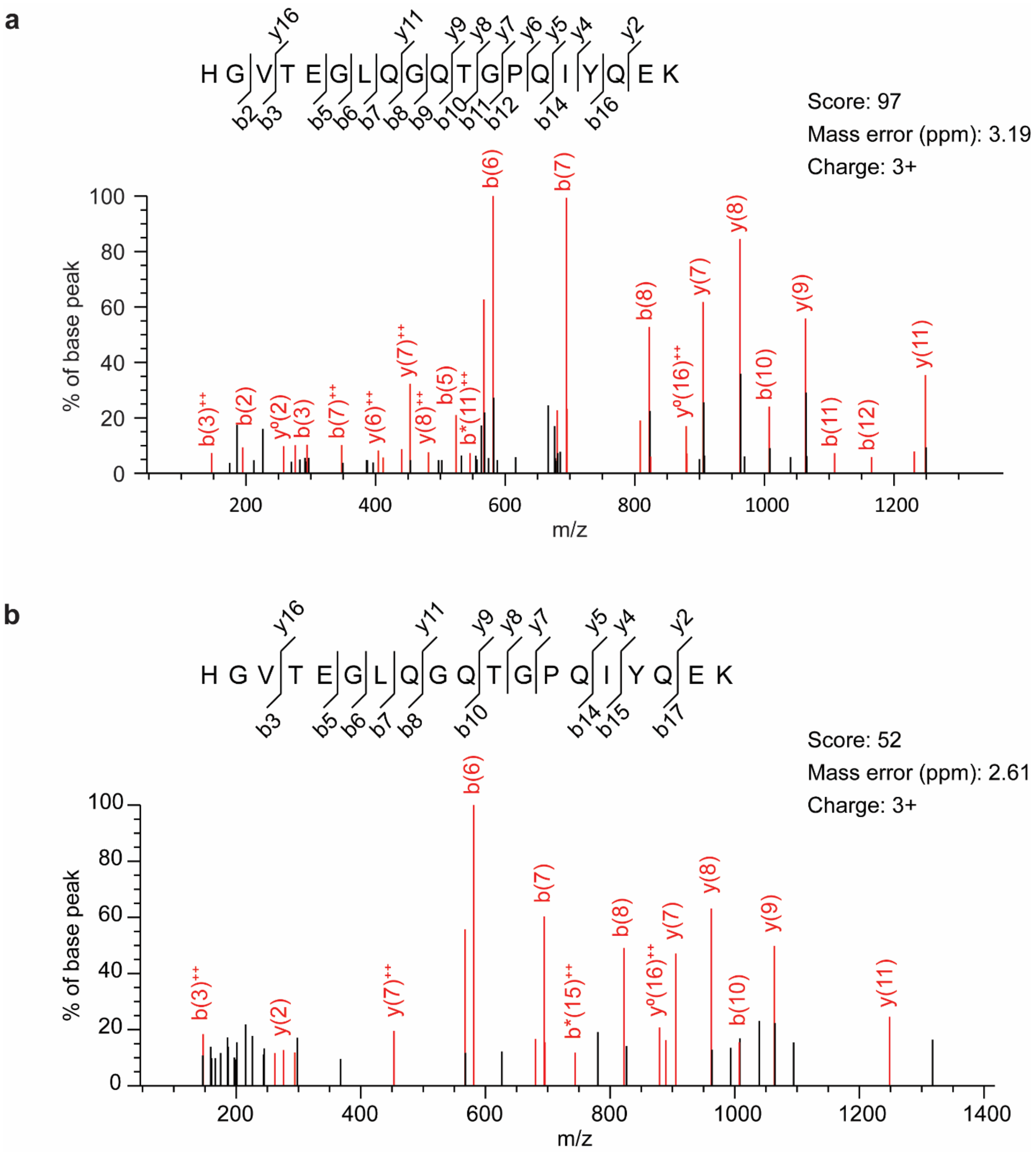


**Supplementary Fig. 1 Alt-RPL36 is expressed in HT1080 and MOLT4 cells.** MS/MS spectra of a unique alt-RPL36 tryptic peptide detected via peptidomics in HT1080 cells (**a**) and MOLT4 cells (**b**).

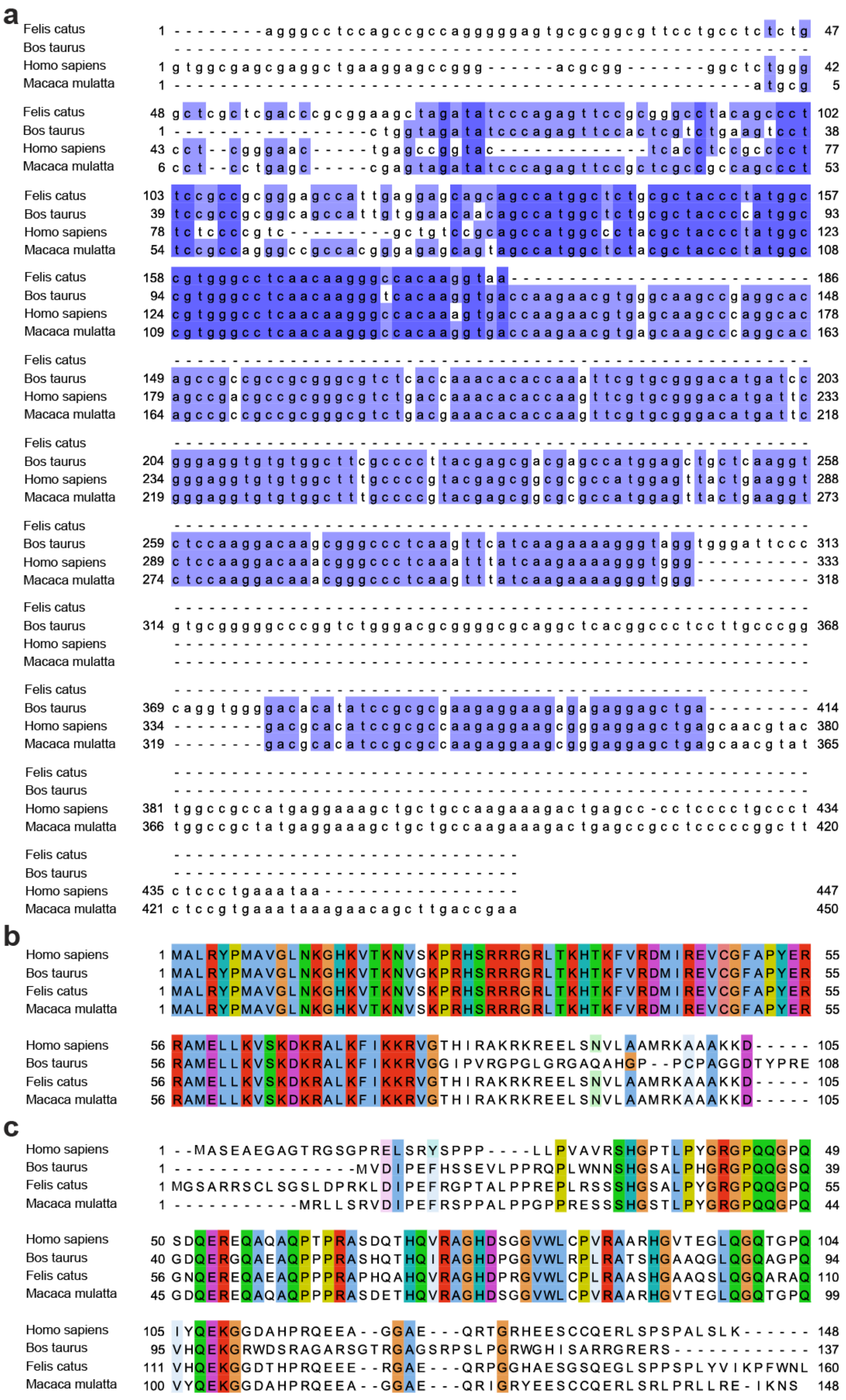
a
Felis catus
Bos taurus
Homo sapiens
Macaca mulatta
b
Homo sapiens
Bos taurus
Felis catus
Macaca mulatta
c
Homo sapiens
Bos taurus
Felis catus
Macaca mulatta

**Supplementary Fig. 2 Alt-RPL36 is predicted to be conserved in some mammals. a** mRNA sequence conservation of RPL36 transcript isoforms in *Bos taurus* (XR_003035408.1), *Felis catus* (XM_011287701.3), and *Macaca mulatta* (XM_015122577.2) at the nucleotide level. Conserved nucleotides are highlighted in blue. Dark blue indicates identity, while light blue indicates conservation in at least 3 of 4 species. Species with cognate or near-cognate start codons in strong Kozak consensus sequence in the alt-RPL36 reading frame were chosen for analysis. **b-c** ClustalW2 alignment of RPL36 (**b**) and alt-RPL36 (**c**) protein sequences translated from transcripts shown in (**a**), residues are highlighted to indicate identity. ClustalW2 alignments were visualized using JalView.

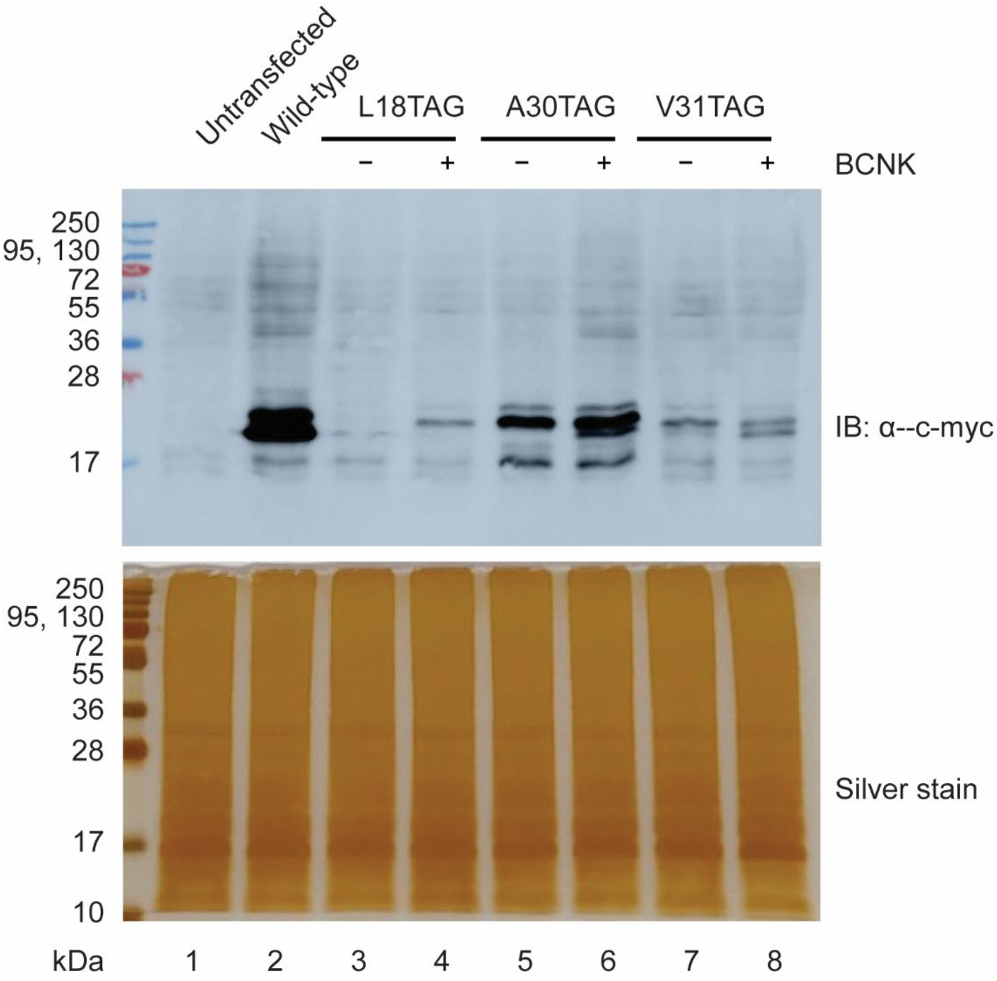


**Supplementary Fig. 3 Expression of alt-RPL36 variants via genetic code expansion.** HEK 293 cells were transfected with transgenes encoding *M. mazei* pyrrolysyl-tRNA synthetase bearing Y306A/Y384F mutations (PylRS-AF), amber-suppressing Pyl tRNA (PylTCUA), and variants of amber codon (TAG)-bearing, c-myc-tagged alt-RPL36. Cells were incubated in the presence or absence of 60 µM bicyclononyne-lysine (BCNK) for 45 hours at 37℃, lysed, and analyzed by Western blotting with anti-c-myc antibody (upper), silver stain served as a loading control (lower). Controls with untransfected cells (lane 1) and cells expressing wild-type alt-RPL36 (lane 2) are shown.

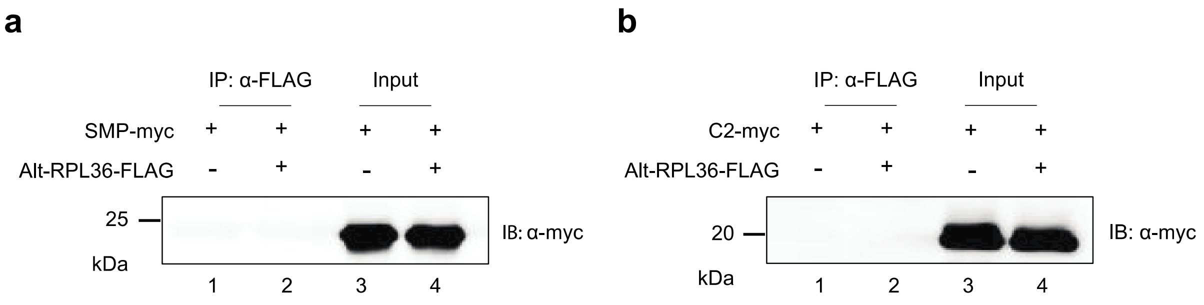


**Supplementary Fig. 4 Both the SMP and the C2 domain are required for the interaction of TMEM24 with alt-RPL36. a** and **b** Control HEK 293T cells (lane 1) or HEK 293T cells stably expressing alt-RPL36-FLAG-HA (lane 2) were transfected with the indicated TMEM24 truncation mutants, and immunoprecipitation (IP) was performed with anti-Flag antibody, followed by immunoblotting (IB) with anti-myc antibody. Cell lysates (1%) before IP (input, lanes 3 and 4) were used as the loading controls. Data are representative of three biological replicates.

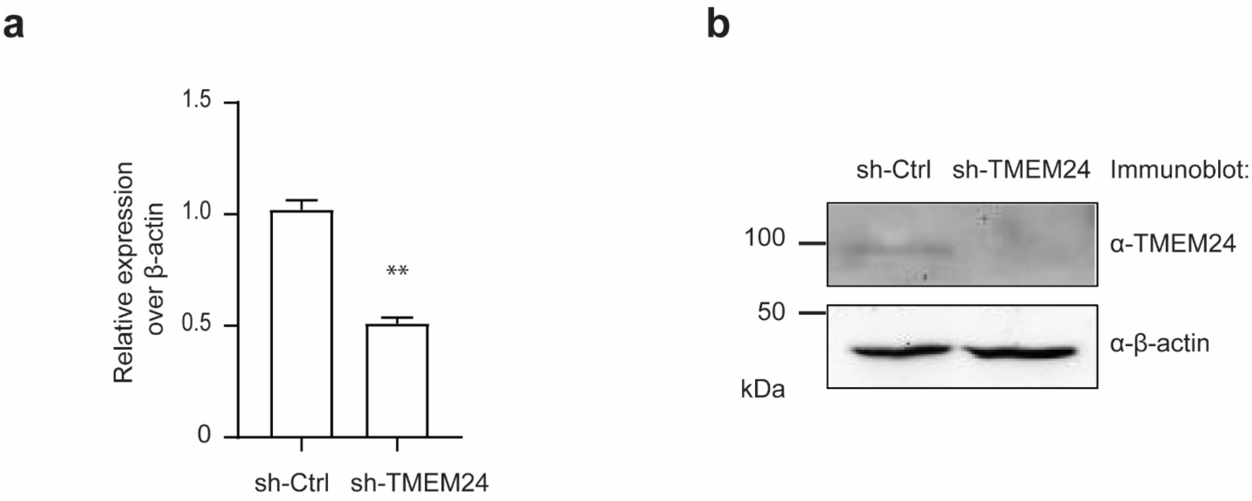


**Supplementary Fig. 5 Silencing TMEM24 in alt-RPL36-FLAG knock-in (KI) cells. a** Quantitative RT-PCR of alt-RPL36 KI cells stably expressing control shRNA (sh-Ctrl) or TMEM24 shRNA (sh-TMEM24) with primer specific to TMEM24 (error bars, standard error of the mean (s.e.m.), $N = 3$, ** $p < 0.01$ (two-tailed $t$-test)). **b** Western blot analysis of alt-RPL36 KI cells stably expressing control shRNA (sh-Ctrl) or TMEM24 shRNA (sh-TMEM24) with the indicated antibodies. Data are representative of two biological replicates.

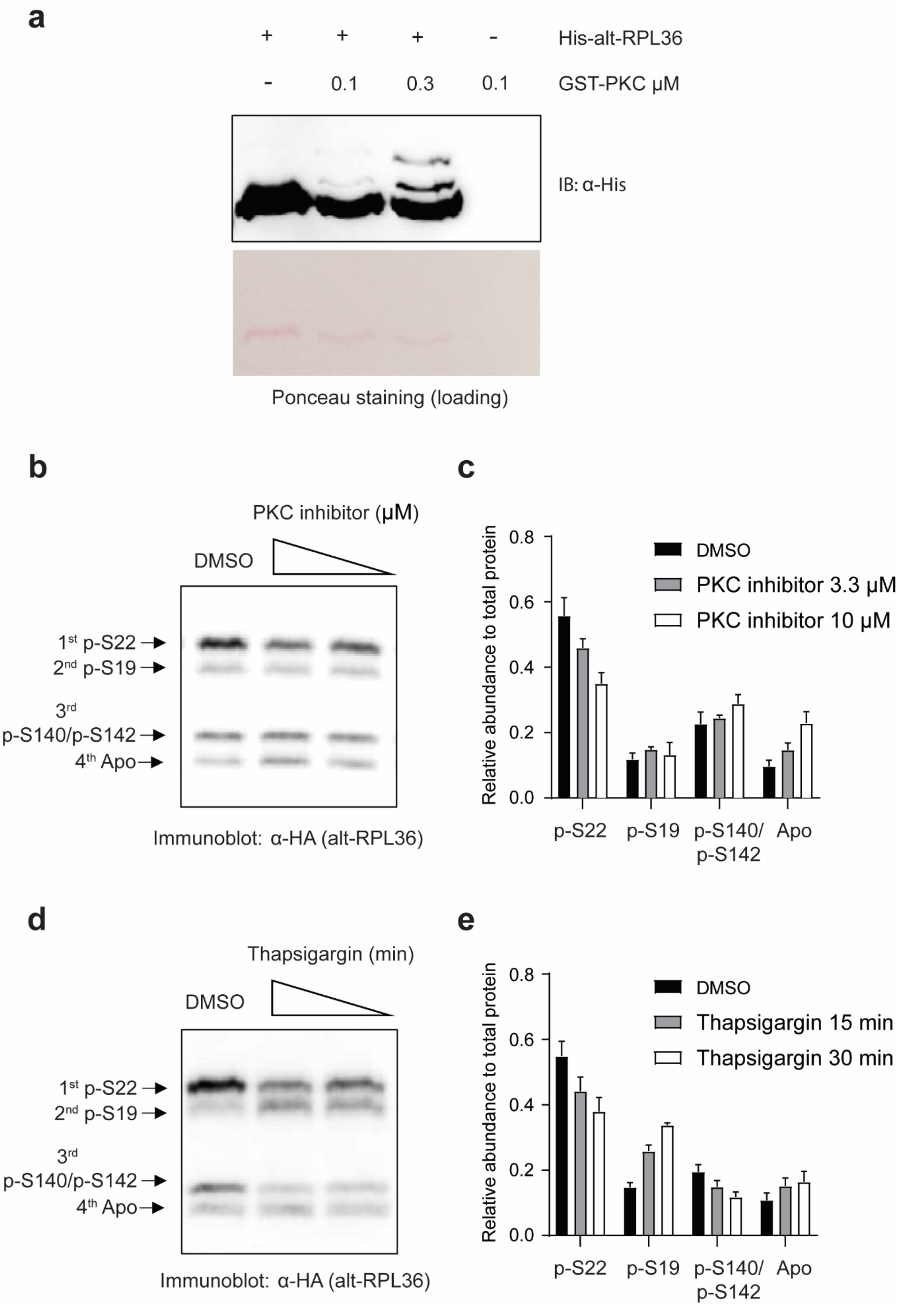
a
+ + + - His-alt-RPL36
- 0.1 0.3 0.1 GST-PKC μM
IB: α-His
Ponceau staining (loading)
b
PKC inhibitor (μM)
DMSO
1st p-S22
2nd p-S19
3rd p-S140/p-S142
4th Apo
Immunoblot: α-HA (alt-RPL36)
c
Relative abundance to total protein
0.0
0.2
0.4
0.6
0.8
DMSO
PKC inhibitor 3.3 μM
PKC inhibitor 10 μM
p-S22
p-S19
p-S140/ p-S142
Apo
d
Thapsigargin (min)
DMSO
1st p-S22
2nd p-S19
3rd p-S140/p-S142
4th Apo
Immunoblot: α-HA (alt-RPL36)
e
Relative abundance to total protein
0.0
0.2
0.4
0.6
0.8
DMSO
Thapsigargin 15 min
Thapsigargin 30 min
p-S22
p-S19
p-S140/ p-S142
Apo

**Supplementary Fig. 6 Phosphorylation of alt-RPL36 is regulated by protein kinase C (PKC) and cytosolic calcium level. a** Purified alt-RPL36 was allowed to react with increasing amounts of PKC, and the reaction was analyzed by Phos-tag Western blotting with anti-His tag antibody. Ponceau staining served as a loading control. **b** HEK 293T cells stably expressing alt-RPL36-FLAG-HA were treated with increasing amounts of PKC inhibitor or vehicle for 8 hr, followed with Phos-tag Western blotting. **c** Quantitative analysis of the Western blot signal of the bands in (**b**). Data represent mean values ± standard error of the mean (s.e.m.) of three biological replicates. **d** HEK 293T cells stably expressing alt-RPL36-FLAG-HA were treated with 1 μM thapsigargin for different times or vehicle, followed with Phos-tag Western blotting. **e** Quantitative analysis of the Western blot signal of the bands in (**d**). Data represent mean values ± standard error of the mean (s.e.m.) of three biological replicates.

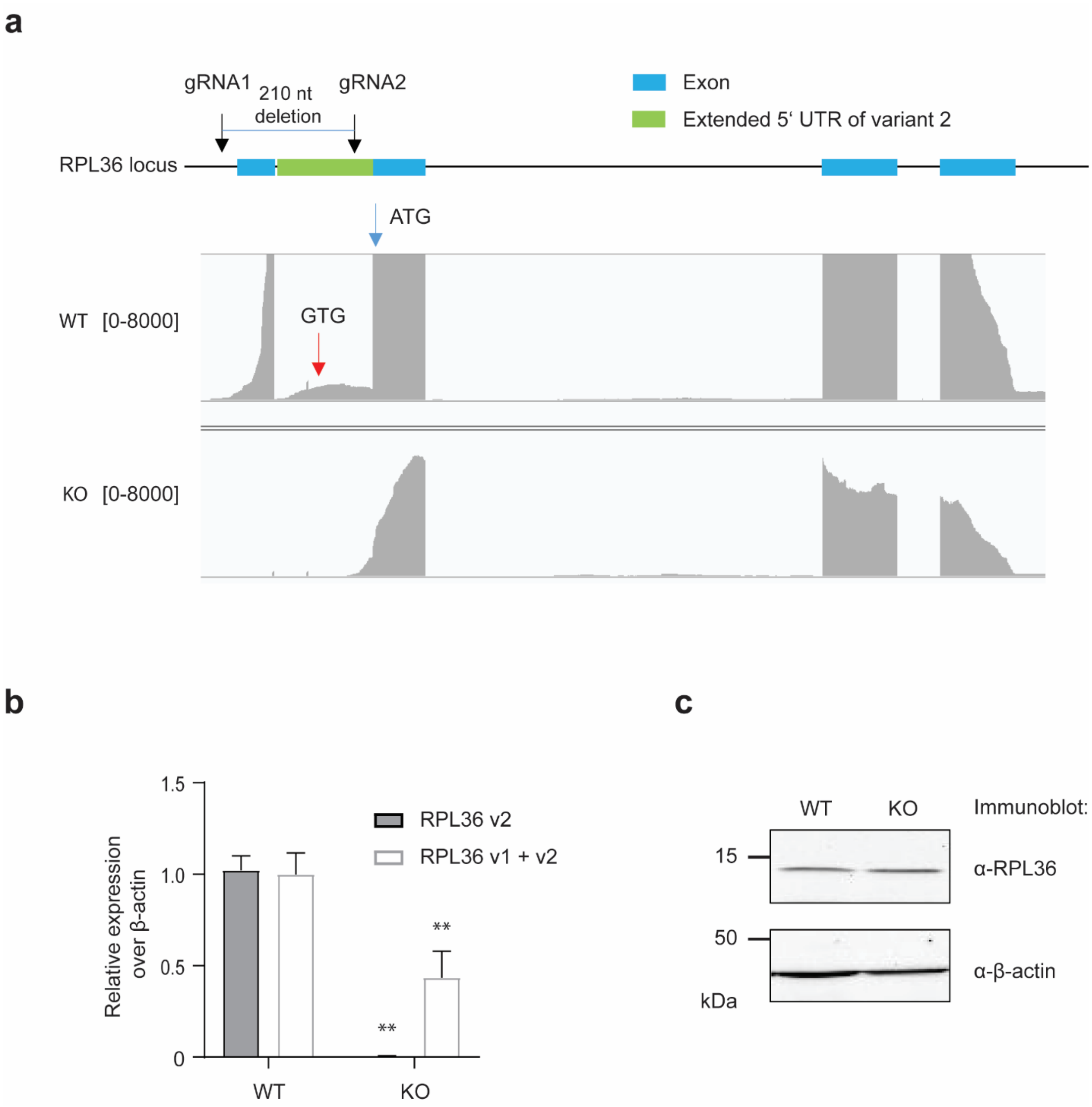


**Supplementary Fig. 7 Validation of the alt-RPL36 knock-out HEK 293T cell line. a** Top, schematic representation of the human *RPL36* genomic locus. The first (blue) exon is specific to transcript variant 1; the second (green) exon is specific to transcript variant 2 and encodes the alt-RPL36 start codon; downstream blue exons are shared by both transcript variants and encode the annotated RPL36 coding sequence. Bottom, RNA-seq reads mapping to the *RPL36* genomic locus in wild-type (WT) and alt-RPL36 knock-out (KO) cells. The position of the GTG start codon of alt-RPL36 is indicated with a red arrow, and the position of the ATG start codon of RPL36 is indicated with a blue arrow. The y-axis scale for RNA is reads per kilobase per million reads (y-axis scale, 0-8000 RPKM). **b** Quantitative RT-PCR with primers specific to RPL36 variant 2 (RPL36 v2) or primers that detect both variants (RPL36 v1 and v2) (error bars, standard error of the mean (s.e.m.), $N = 6$, ** $p < 0.01$ (two-tailed *t*-test)). **c** Western blot analysis of the wild-type and alt-RPL36 knock-out HEK 293T cells with the indicated antibodies. Data are representative of three biological replicates.

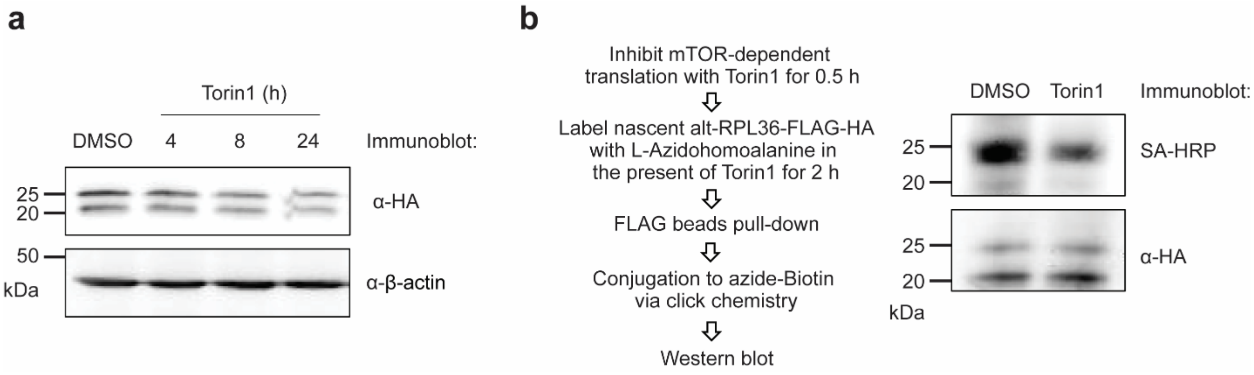


**Supplementary Fig. 8 mTOR inhibition downregulates the translation of alt-RPL36. a** HEK 293T cells stably expressing alt-RPL36-FLAG-HA were treated with 0.5 μM mTOR inhibitor Torin1 for different times or vehicle, followed with Western blot. **b** Left: experimental scheme for newly translated alt-RPL36-FLAG-HA capture. Right: anti-HA reflects the total protein level of alt-RPL36-FLAG-HA, and streptavidin-HRP (SA-HRP) reflects newly synthesized alt-RPL36-FLAG-HA.

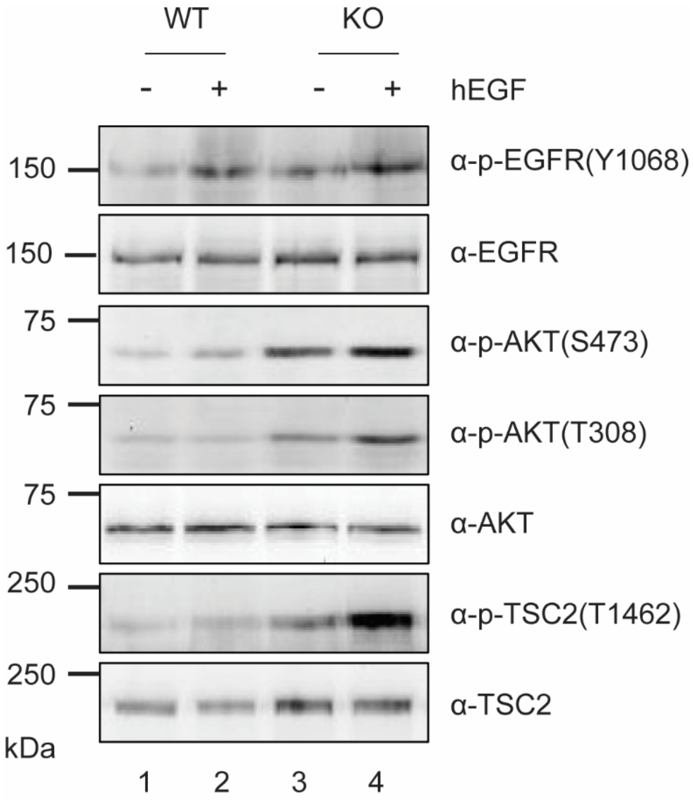


**Supplementary Fig. 9 Alt-RPL36 knock-out cells are more sensitive to EGF stimulation, compared to wild-type cells.** Wild-type (WT) or alt-RPL36 knock-out (KO) HEK 293T cells were treated with FBS-free media for 6 h, followed by 100 ng/mL hEGF stimulation for 15 min or control (no stimulation). Cells were collected, and Western blot performed with the antibodies indicated on the right. All Western blots are representative of two biological replicates.

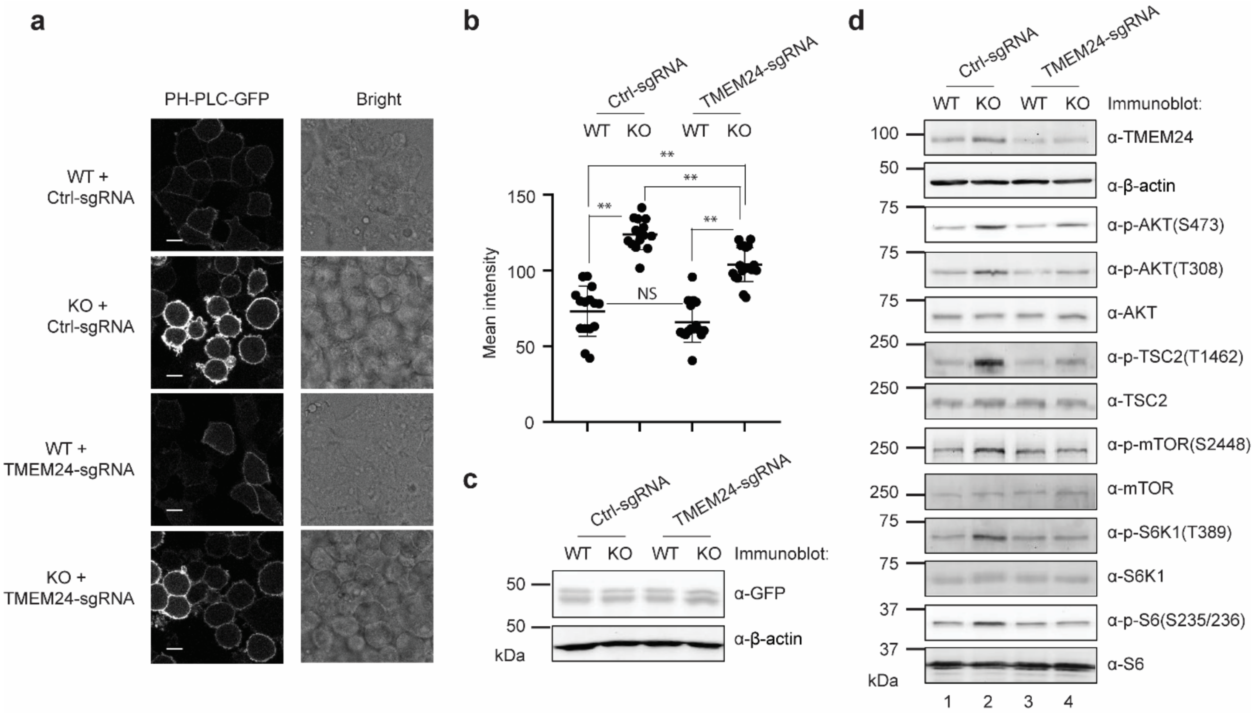


**Supplementary Fig. 10 Regulation of PI3K-AKT-mTOR pathway by alt-RPL36 requires its interaction with TMEM24. a** PH-PLC-GFP reporter was stably introduced into the following cell lines: HEK 293T cells stably expressing Cas9 and control sgRNA (WT + Ctrl-sgRNA), alt-RPL36 KO cells stably expressing Cas9 and control sgRNA (KO + Ctrl-sgRNA), HEK 293T stably expressing Cas9 and TMEM24 sgRNA (WT + TMEM24-sgRNA), alt-RPL36 KO stably expressing Cas9 and TMEM24 sgRNA (KO + TMEM24-sgRNA). Confocal imaging was performed with live cells. Scale bar, 10 µm. **b** Quantitation of the PH-PLC-GFP signals in the four cell lines described above. At least 20 fields of view were analyzed, totaling > 400 cells for each measurement. Significance was evaluated with two-tailed *t*-test. ** $p < 0.01$. NS, not significant. **c** Western blot of the four cell lines with antibodies indicated on the right for comparison of PH-PLC-GFP expression. Data are representative of three biological replicates. **d** Western blot analysis of the four cell lines described above with antibodies indicated on the right. Data are representative of three biological replicates.

**a**

| Mutation data obtained from COSMIC | | | |
|---|---|---|---|
| CDS mutation | RPL36 AA mutation | alt-RPL36 AA mutation | Primary Tissue |
| 49G>A | V17M | S50N | Skin |
| 194A>C | K65T | Q98H | Large intestine |
| 212A>C | K71T | Q104H | Urinary tract |
| 243C>G | I81M | P115A | Breast |
| 254G>A | R85K | Synonymous | Large intestine |
| 284C>T | A95V | Synonymous | Stomach |
| 117C>T | Synonymous | R73C | Upper aerodigestive tract |
| 117C>T | Synonymous | R73C | Urinary tract |
| 178T>C | Synonymous | V93A | Liver |
| 252G>A | Synonymous | E118K | Cervix |

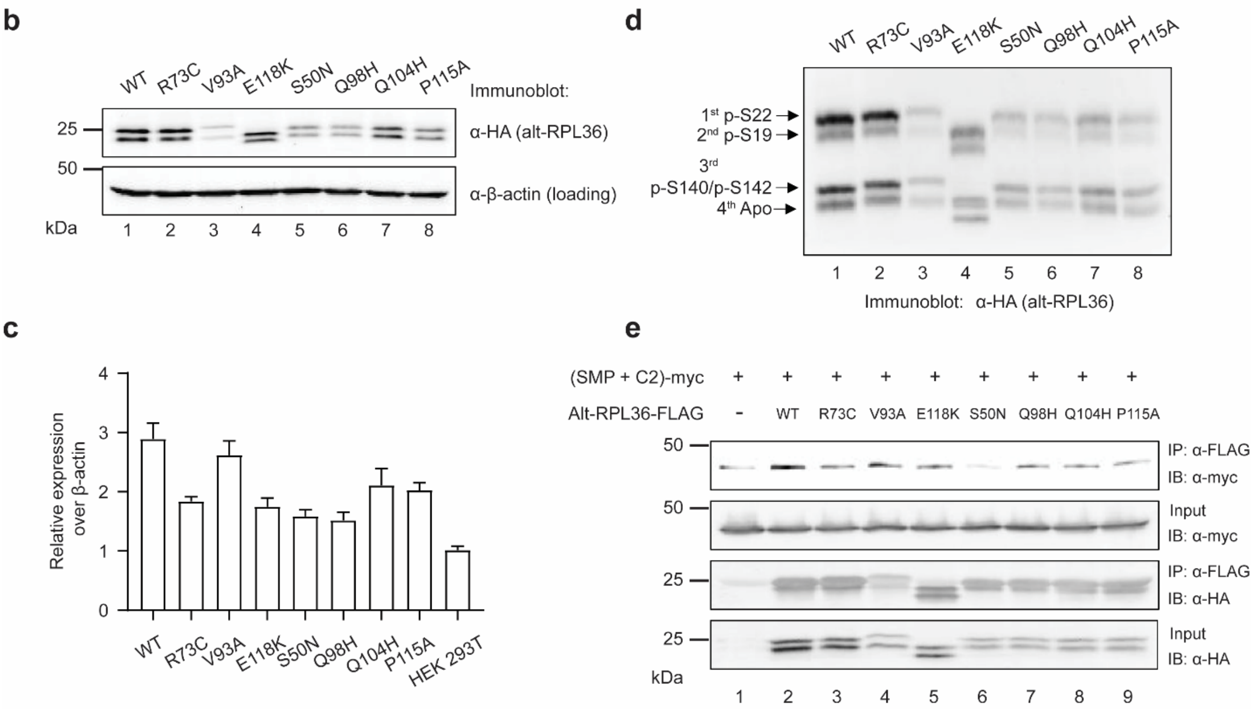


**Supplementary Fig. 11 Cancer-related mutations affect alt-RPL36 protein properties and interactions. a** Cancer-associated non-synonymous mutations in *RPL36* and/or *alt-RPL36* from the COSMIC database. AA, amino acid. COSMIC DNA numbering is relative to the *RPL36* start codon ($A_1$TG). **b** SDS-PAGE Western blot of HEK 293T cells stably expressing wild-type (WT) alt-RPL36 or point mutants indicated at the top with antibodies indicated on the right. **c** Quantitative RT-PCR of control HEK 293T cells (HEK 293T) or the stable cell lines indicated at the bottom using primers designed against *RPL36* transcript variant 2 that are specific to the alt-RPL36 coding sequence (error bars, standard error of the mean (s.e.m.), $N$ = 3, ** $p$ < 0.01 (two-tailed $t$-test)). **d** Wild-type alt-RPL36-FLAG-HA or point mutants indicated at the top were immunopurified from HEK 293T cells, resolved with Phos-tag SDS-PAGE, and detected with anti-HA via Western blotting. **e** Control HEK 293T cells (lane 1), HEK 293T cells stably expressing wild-type alt-RPL36-FLAG-HA (lane 2), and HEK 293T cells stably expressing point mutants indicated at the top (lane 3 to 9) were transfected with the TMEM24 SMP + C2 domain construct, followed by IP with anti-FLAG antibody and immunoblotting with antibodies indicated on the right. Cell lysates (1%) before IP (input) were used as loading controls. All Western blots are representative of two biological replicates.

a

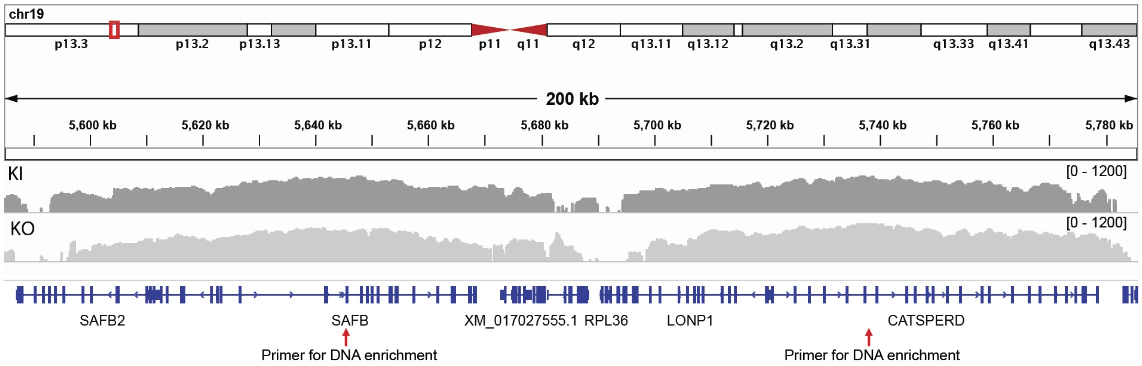


b

| Chromosome | Position | Cell Line | Type of variant | Supporting Reads | Gene | Intron/Exon |
|---|---|---|---|---|---|---|
| chr19 | 5708507 | KO | Insertion | 125 | LONP1 | Intron |
| chr19 | 5740515-5742010 | KO | Inversion | 20 | CATSPERD | Intron |
| chr19 | 5741045-5741079 | KO | Deletion | 376 | CATSPERD | Intron |
| chr19 | 5761620 | KO | Insertion | 15 | CATSPERD | Intron |
| chr19 | 5708507 | KI | Insertion | 81 | LONP1 | Intron |
| chr19 | 5741045-5741079 | KI | Deletion | 235 | CATSPERD | Intron |
| chr19 | 5761620 | KI | Insertion | 25 | CATSPERD | Intron |

**Supplementary Fig. 12 No large deletions or insertions were observed in the 100 kb upstream or 100 kb downstream of RPL36 in the alt-RPL36 knock-out and knock-in cell lines.** Shown are the aligned Xdrop sequencing results targeting 100 kb upstream and 100 kb downstream of RPL36 (**a**), visualized via IGV. The primer sites selected for targeted DNA enrichment are indicated with a red arrow. Four structural variants were detected in the targeted region (**b**), three of which are shared by the knock-out and knock-in cell lines, and one inversion specific to the knock-out cell line localized to an intronic sequence within the CATSPERD gene. No cell-line specific insertions or deletions were detected. We did not observe the inserted sequence in the knock-in cells, possibly because of low coverage of the RPL36 gene.

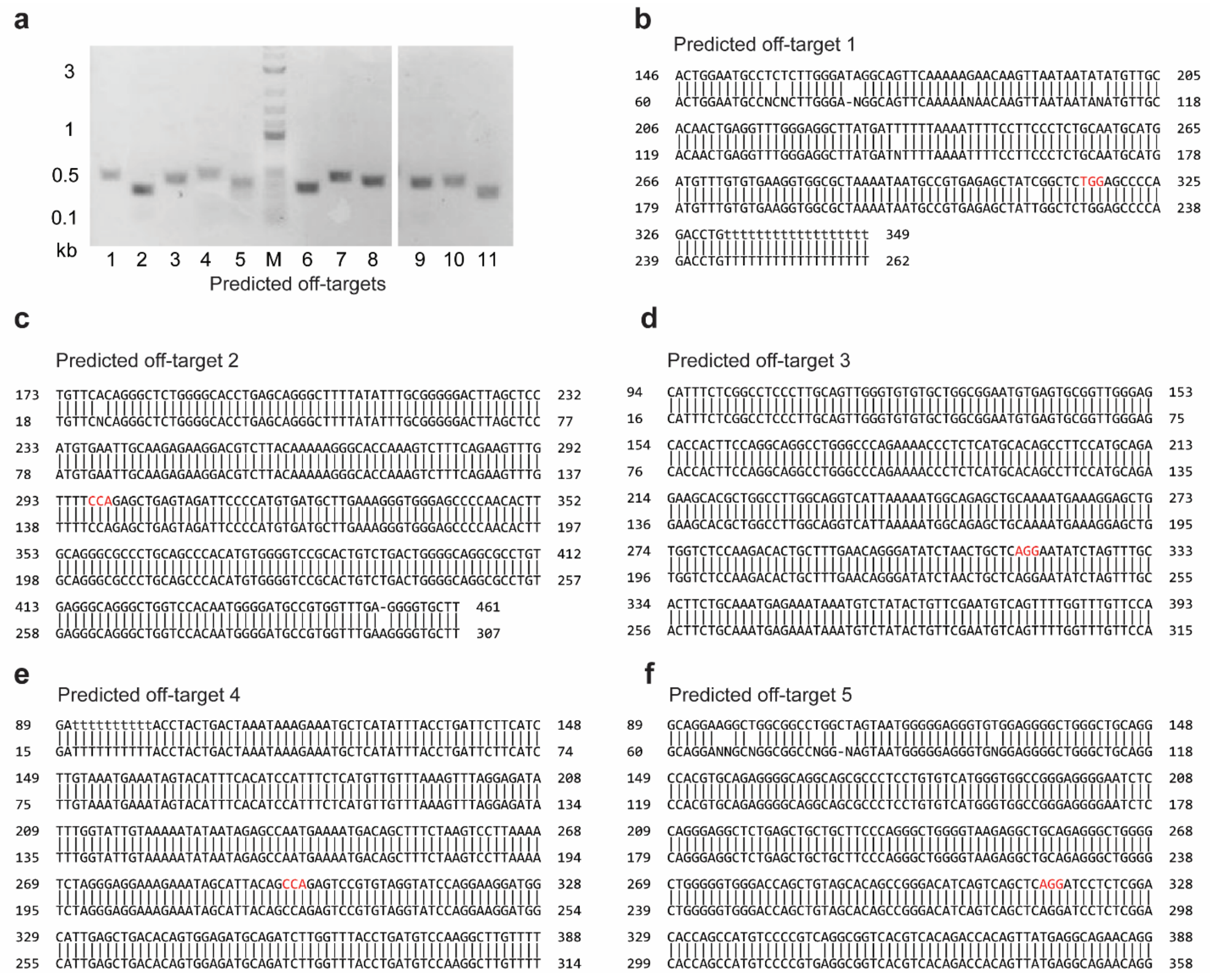
a
3
1
0.5
0.1
kb
1 2 3 4 5 M 6 7 8 9 10 11
Predicted off-targets
b
Predicted off-target 1
c
Predicted off-target 2
d
Predicted off-target 3
e
Predicted off-target 4
f
Predicted off-target 5

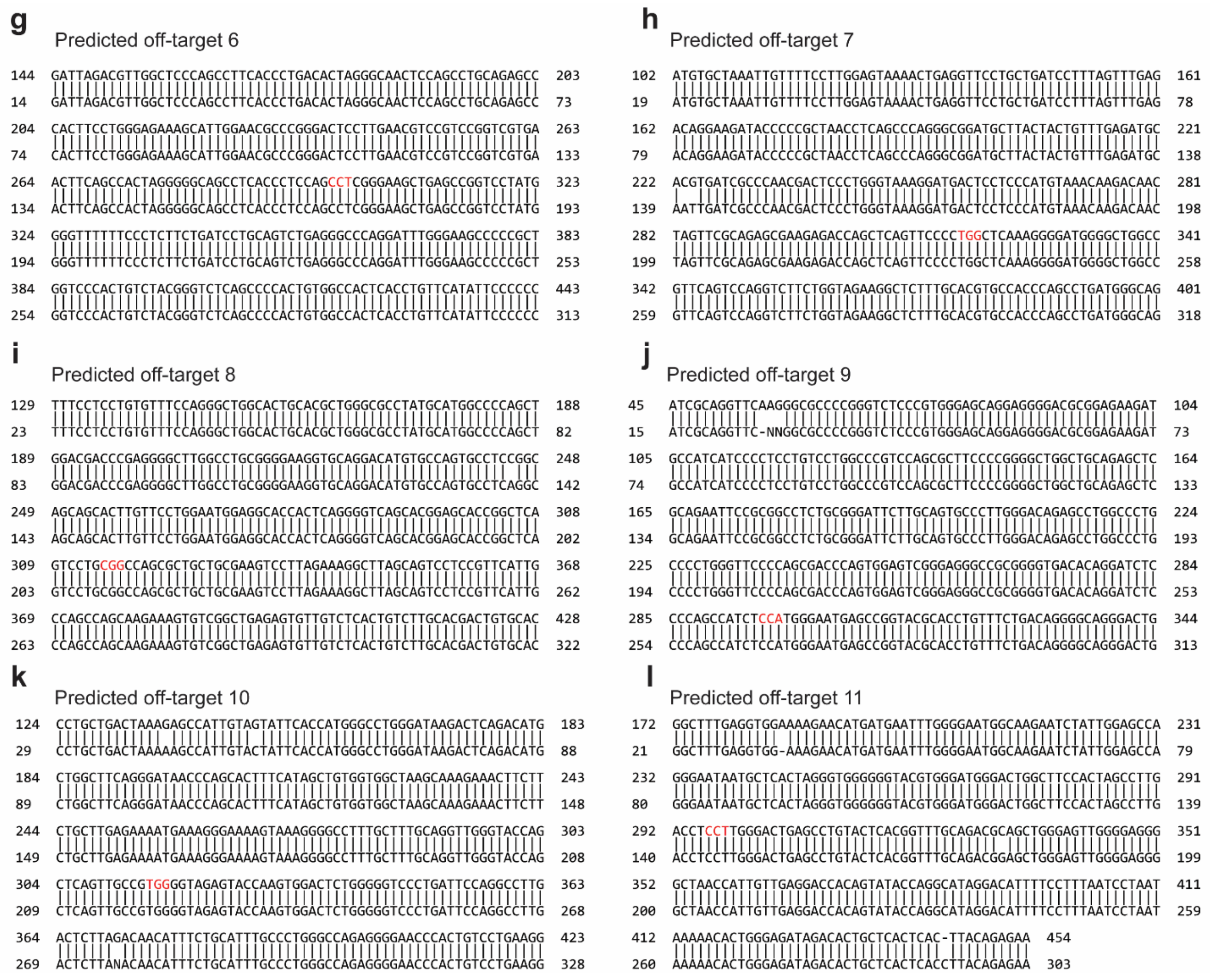


**Supplementary Fig. 13 alt-RPL36 knock-out cells do not contain deletions or insertions at 11 predicted off-target sites. a** Genomic PCR and sequencing of 11 off-target loci predicted by cas-offinder separated via agarose gel electrophoresis. **b-l** Sanger sequencing of 11 gel-purified genomic PCR products, followed with BLAST alignment. The NGG PAM sites of the predicted off-target guide RNA are indicated in red. Predicted off-targets 2, 4, 6, 9 and 11 target the antisense sequence. No mutations are observed near the NGG PAM sites.

| RT-qPCR primers (5' - 3') | | | |
|---|---|---|---|
| Target genes | Use | Forward primer | Reverse primer |
| RPL36 v2 | RT-qPCR | AGGTTGGAGGATGGTTGGTT | TGCTCACGTTCTTGGTCACTT |
| RPL36 v1 + v2 | RT-qPCR | ACTGAAGGTCTCCAAGGAC | TCTTTATTTCAGGGAGAG |
| COL6A2 | RT-qPCR | CTCCTCGGGACCAGGACTT | GGTGTCCAGCACGAAGTACA |
| EFNA3 | RT-qPCR | ACAGCCCCATCAAGTTCTCG | GAGTGGGCGTGGAGATGTAG |
| PPP2R2C | RT-qPCR | CAGCTATGTGACTGAAGCTGAC | TAGTCAAACTCCGGCTCGTG |
| TMEM24 | RT-qPCR | TTCCCAGTCTCTGTGACCCA | GATATGAGCAGCCCCTCACC |
| β-actin | RT-qPCR | AGGCACCAGGGCGTGAT | GCCCACATAGGAATCCTTCTGAC |

**Supplementary Table 6 RT-qPCR primers (5' - 3').**

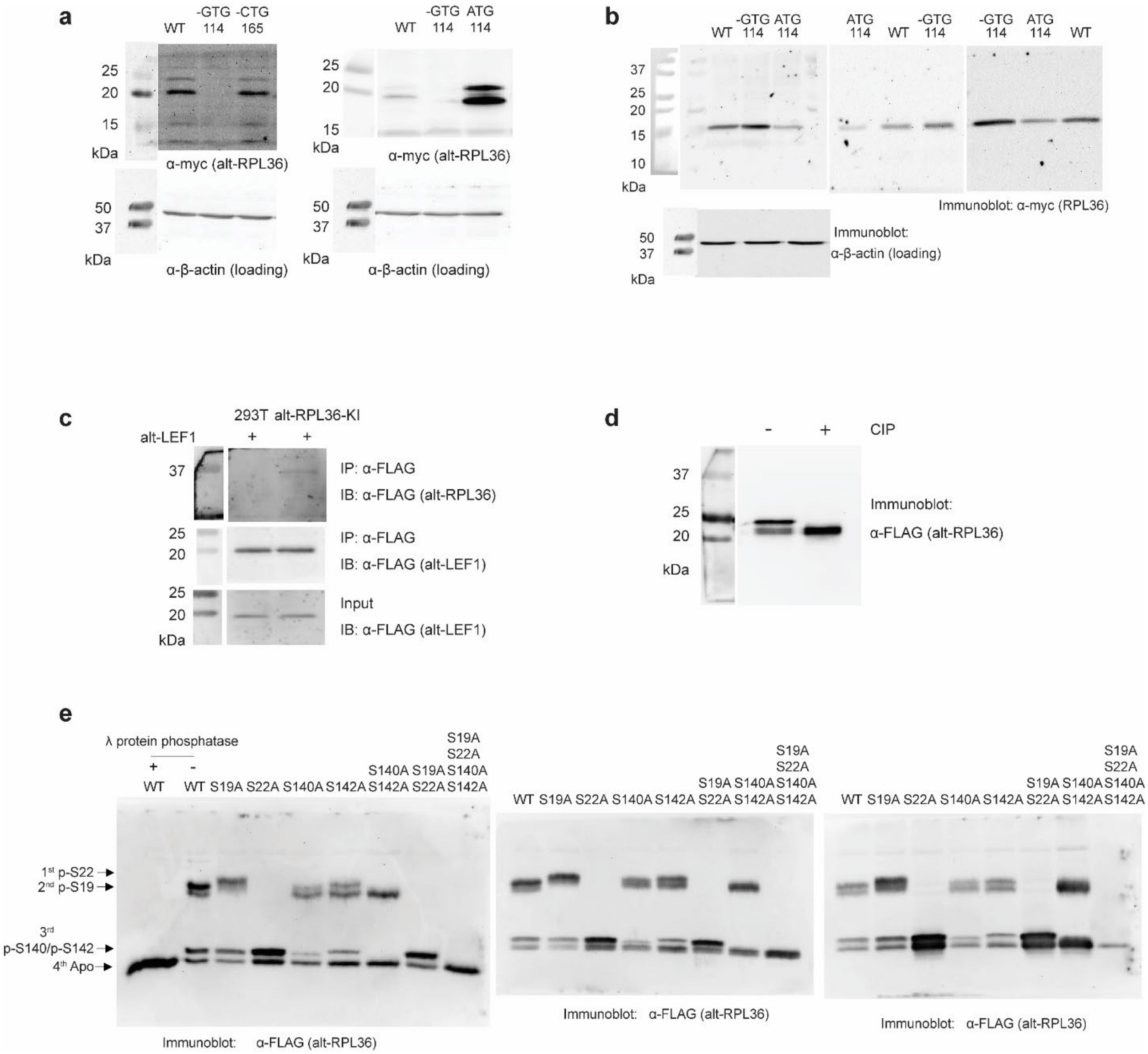


**Supplementary Fig. 14 Uncropped blots from Figure 1, 2 and 3.**

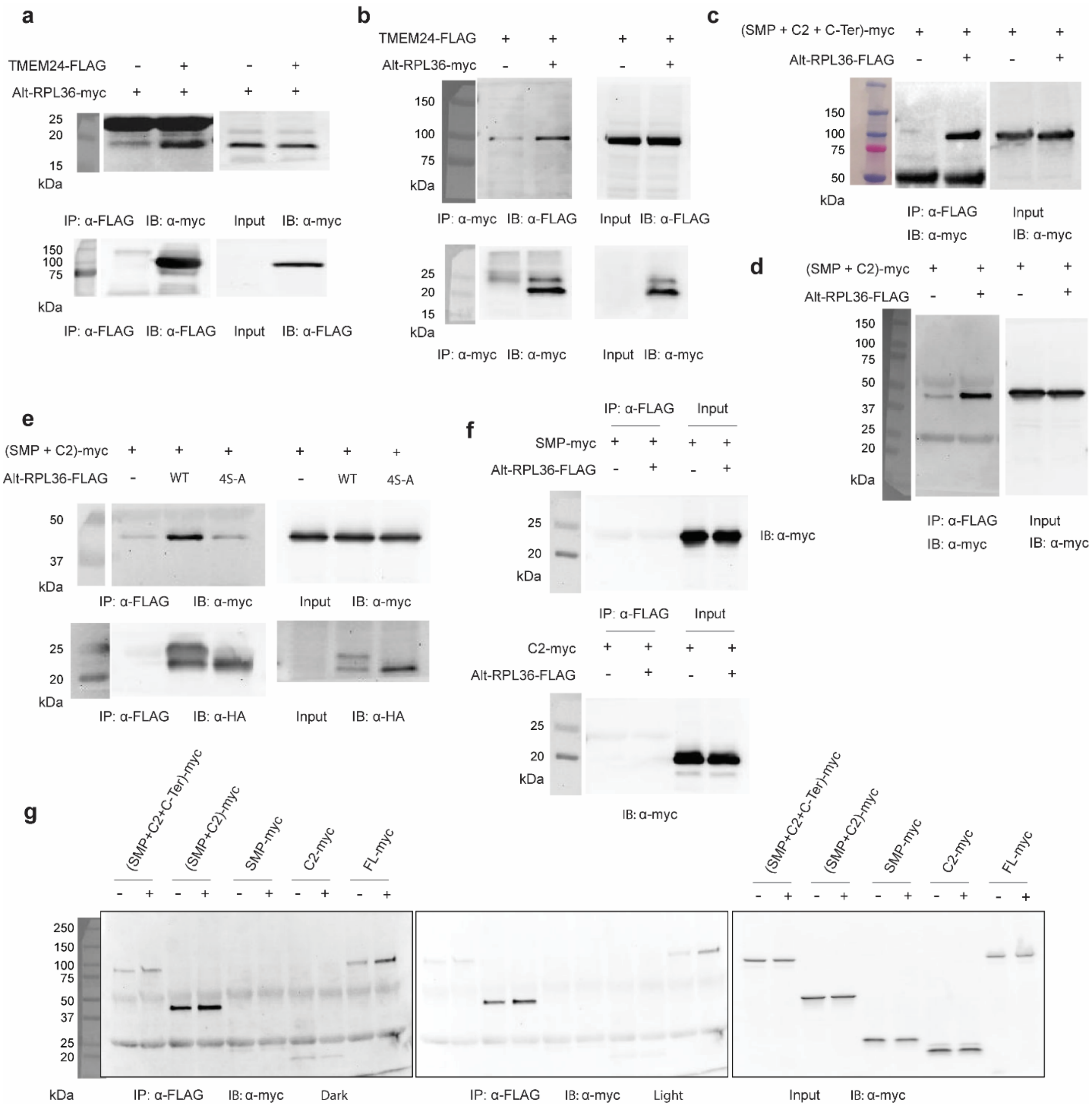


**Supplementary Fig. 15 Uncropped blots from Figure 4 and Supplementary Figure 4.**

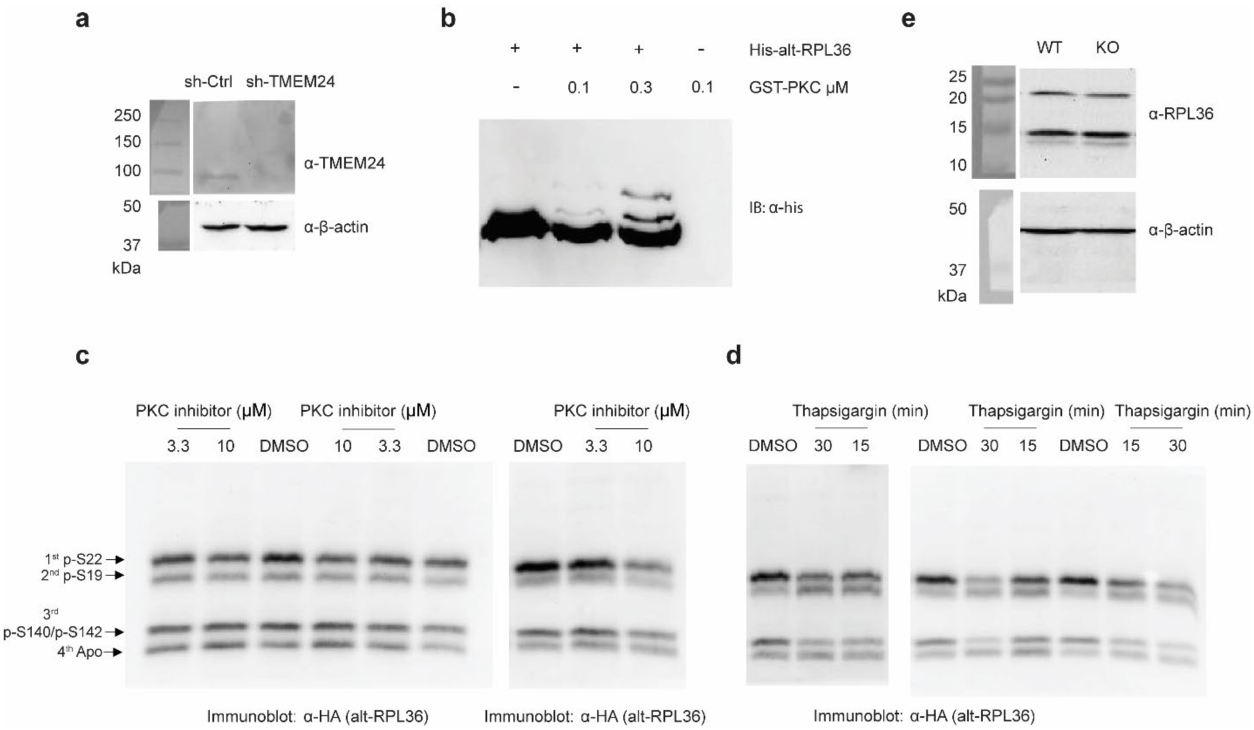


**Supplementary Fig. 16 Uncropped blots from Supplementary Figures 5, 6 and 7.**

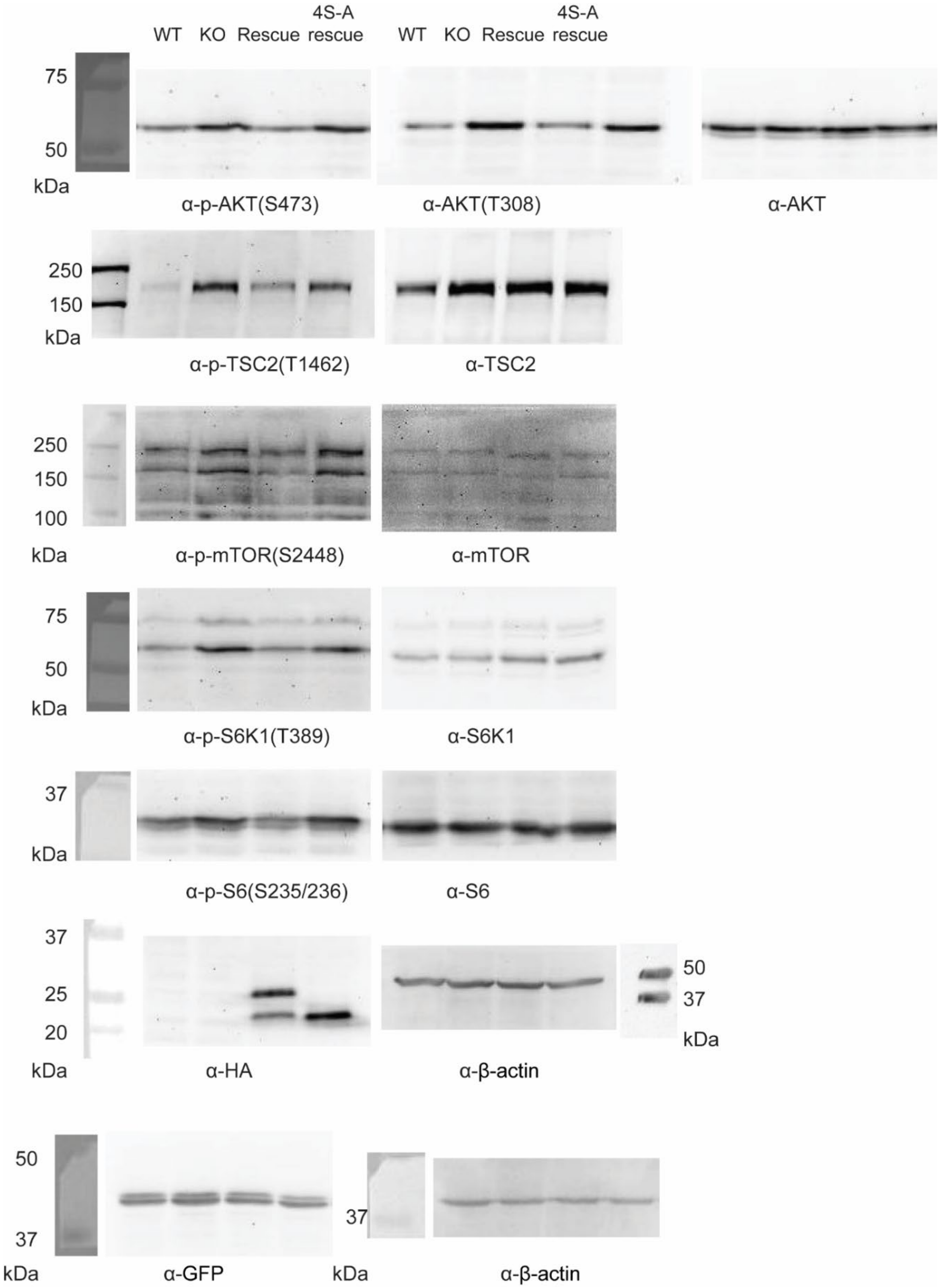


**Supplementary Fig. 17 Uncropped blots from Figure 5.**

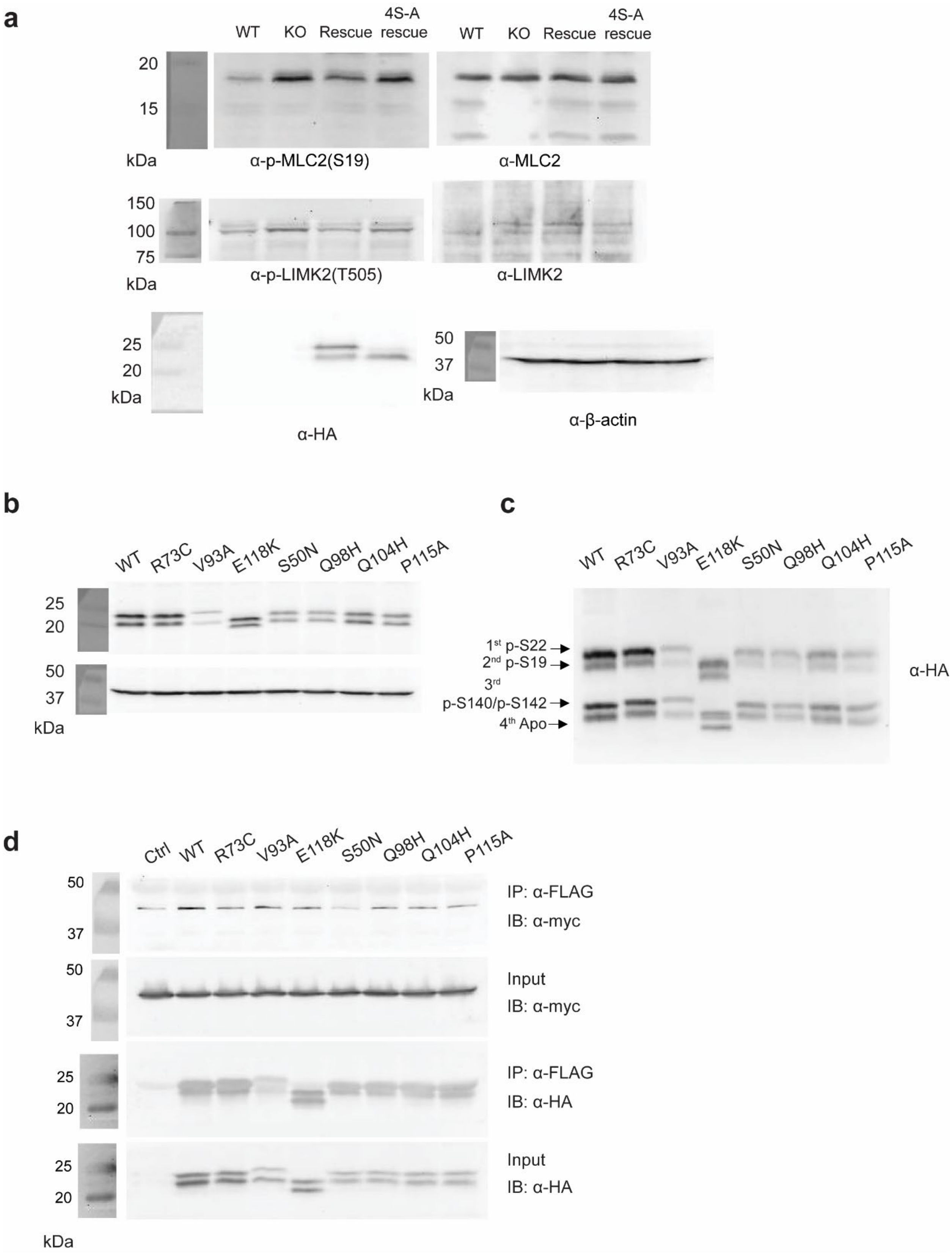


**Supplementary Fig. 18 Uncropped blots from Figure 6 and Supplementary Figure 11.**

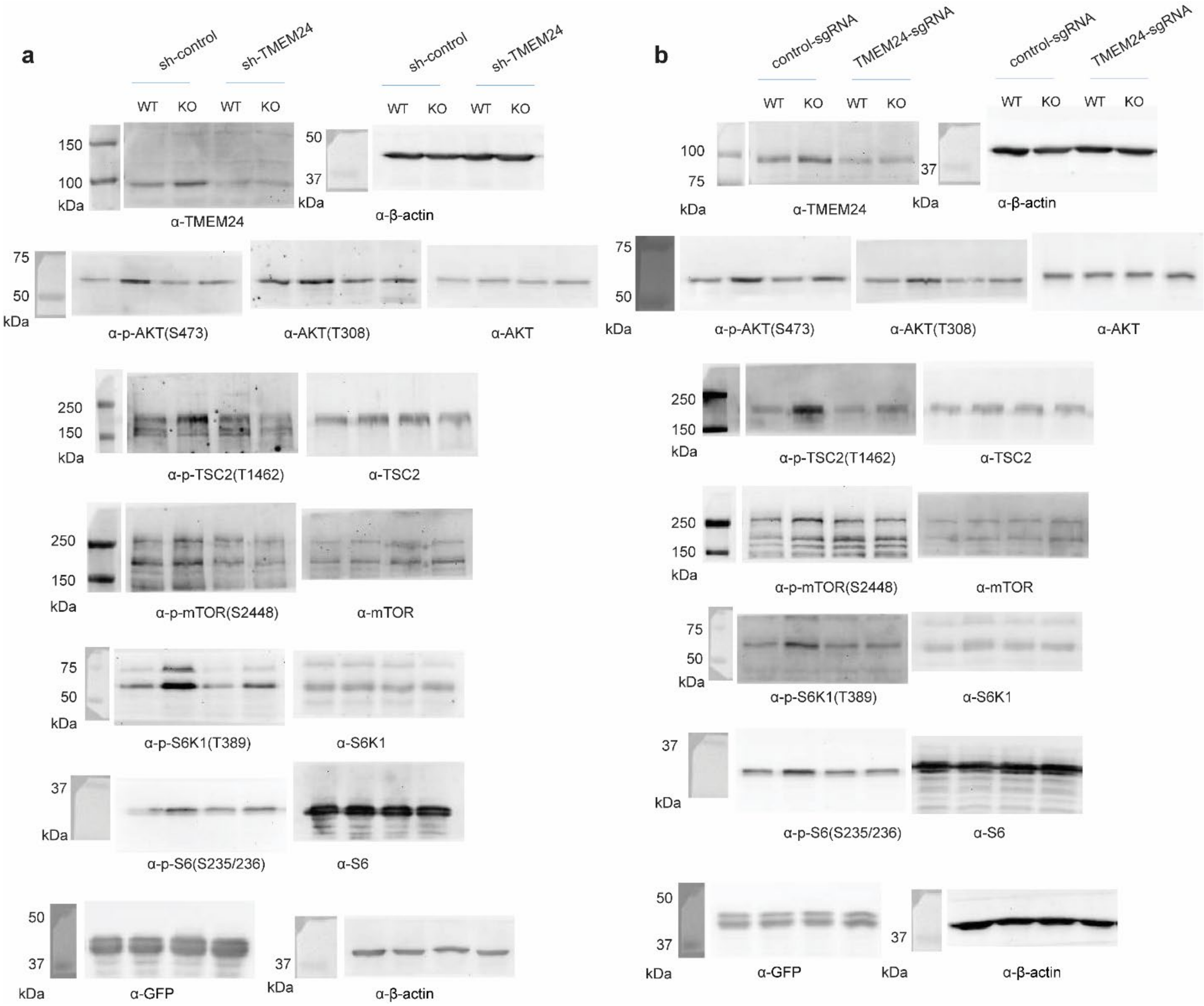


**Supplementary Fig. 19 Uncropped blots from Figure 7 and Supplementary Figure 10.**

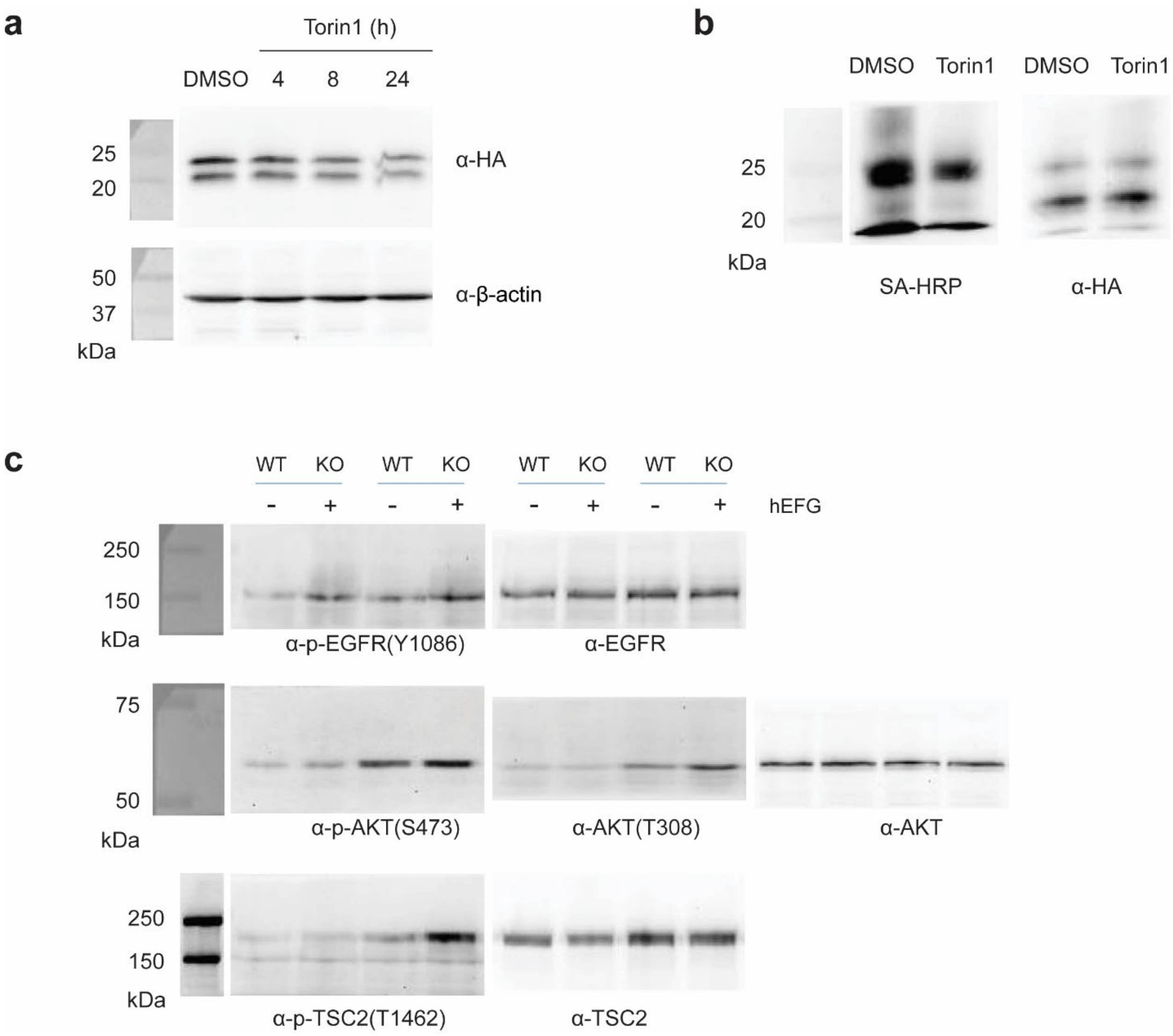


**Supplementary Fig. 20 Uncropped blots from Supplementary Figure 8 and 9.**
